# Inducible Calling Cards: Developing Mouse Reagents for Experimentally Controlled Transposon Insertion *in vivo*

**DOI:** 10.1101/2025.11.04.685928

**Authors:** Simona Sarafinovska, Arthi Venkatesan, Titilope Akinwe, Alexander Chamessian, Xuhua Chen, Maria Payne, Meaghan C. Creed, Robi D. Mitra, Joseph D. Dougherty

## Abstract

The piggyBac transposase enables robust forward genetic screens in mice. Further, fusions of piggyBac with specific transcription factors (TFs) enables recovery of recoverable transposons (Calling Cards) for recording location of DNA binding events *in vitro* and *in vivo*. Such applications would be enhanced by engineering inducible transposases, such that timing of recording could be precisely controlled *in vivo*, as has been previously developed *in vitro.* Here, we tested two approaches for applying inducible Calling Cards (iCC) in the murine brain. We engineered knock-ins of inducible versions of two TFs: Jun, an immediate early gene that serves as a proxy for neural activity, and Sp1, a promiscuous binder of CpG unmethylated regions that indicates active promoters. We fused hyperPiggyBac transposase and a tamoxifen-inducible domain (ERT2) to these TFs and tested the system for efficacy and temporal control of insertions, both *in vitro* and *in vivo*. Jun-iCC mice developed normally with no behavioral abnormalities, showed tamoxifen-dependent recording, and captured neural activity during pharmacologically induced seizures. Jun-iCC yields relatively low numbers of insertions, likely due to the transient expression of Jun. In contrast, Sp1-iCC provided substantially higher insertion numbers, but transgenic animals exhibited developmental abnormalities, including reduced viability, anophthalmia, and reduced body weight, suggesting that ERT2 domains may sequester Sp1 and thus significantly impact development. Nonetheless, these inducible Calling Cards mouse lines enable drug-inducible integration of transposon cargo into specific loci in the living mouse.

**SIGNIFICANCE STATEMENT:** Transposases, like the more widely used recombinases (Cre, Flp, etc.), enable the integration of a variety of DNA sequences into the genome, but without dependence on LoxP sites. However, few mouse reagents exist for transposases. Cre has been adapted to be drug-inducible, as well as activity-regulated (e.g., FOS-TRAP). Thus, additional applications would become available if transposases could also be drug-controlled and activity-dependent. Further, there may be benefits to targeting integration to specific genomic region types (e.g., promotors). Here, we report results testing mouse lines for two drug-inducible transposase fusions to transcription factors (TF) – the promoter binding SP1, and activity dependent Jun. Both allowed genomic integration, with viability, efficiency, inducibility and targeting varying by TF-fusion.

## INTRODUCTION

Transposons have wide applications for genomics and synthetic biology. PiggyBac, the cut-and-paste transposase derived from *Trichoplusia ni*, excises transposons with inverted terminal repeats and inserts them, and up to 15 kb of DNA cargo, into DNA. It requires only long terminal repeats within the donor transposon and can insert into any available TTAA sequence (Cadiñanos & Bradley, 2007; Troyanovsky et al., 2016). PiggyBac has been further engineered to increase transposition activity, leading to hyperpiggyBac (hyPB) (Cadiñanos & Bradley, 2007). PiggyBac and derivatives have been utilized in mammalian systems for the insertion of expression constructs to create cell lines, and the insertion of gene traps for genetic screens (Chew et al., 2011; Rad et al., 2010; Troyanovsky et al., 2016). For instance, by crossing mouse lines with hyperpiggyBac engineered into the Rosa26 locus with mouse lines carrying arrays of transposons, novel tumor suppressors were identified in a large-scale forward genetic screen (Rad et al., 2010).

Notably, these transposases have affinity for the BRD4 chromatin modifier, which localizes to enhancers. Thus, we have used engineered transposons (‘Calling Cards’), recovered by high-throughput sequencing, to map hyPB activity and, thus, BRD4 binding (Cammack et al., 2020; Moudgil et al., 2020; Yen et al., 2023). Furthermore, fusions of hyPB to transcription factors (TFs) can redirect transposase activity to other loci. This enables an alternative to Cut-N-Run or ChIP-Seq for assessing the binding pattern of specific TFs in the brain, with one additional, unique feature – the Calling Card is a permanent genomic mark of transient DNA binding events. Thus, Calling Cards have potential applications for associating historical epigenetic events with later cellular outcomes (Boros et al., 2025; Witchley et al., 2021).

One drawback of existing CC tools, however, is that once delivered to the brain, they continuously record until sacrifice of the animal; there is no ability to more precisely control the timing of CC activity. To overcome this limitation, we and others previously developed inducible Calling Cards (iCC), using standard drug-controllable systems (e.g., tamoxifen) in culture (Qi et al., 2017). Here, we test two mouse lines for iCC *in vivo*. We utilize a tamoxifen-inducible domain, ERT2(Donocoff et al., 2020), to confer inducibility to Calling Cards recording, engineering hyPB-ERT2 fusions of two TF loci in the genome. Cytoplasmic heat shock protein 90 (HSP90) binds to the ERT2 domain, sequestering the fusion protein to the cytoplasm (Qi et al., 2017). The presence of tamoxifen abolishes HSP90 binding, which in turn allows the fusion protein to enter the nucleus, and thus “turns on” Calling Cards recording (**Figure 1**).

**Figure 1.**
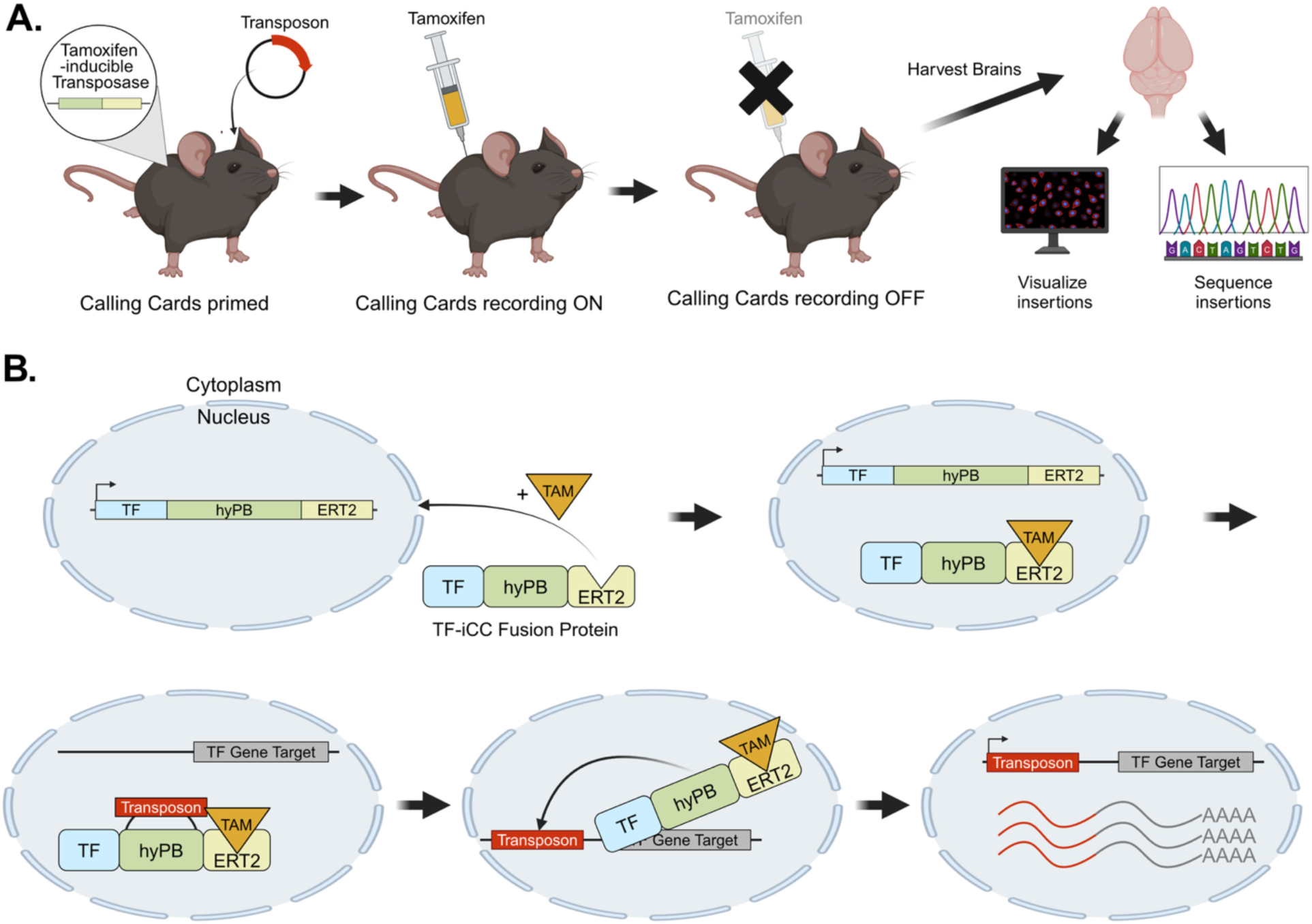
inducible Calling Cards would allow temporally controlled recording of transient molecular states. **A.** The iCC system contains two components: a mouse line with a tamoxifen-inducible transposase knocked into an endogenous locus, and a virally delivered transposon. Presence of these two components “primes” Calling Cards for recording of transposon insertions to turn ON upon systemic delivery of tamoxifen (or its metabolites) for the desired amount of time. Calling Cards recording “turns OFF” with clearance of tamoxifen from the system. The recorded transposon insertions can be visualized using immunofluorescence or sequenced via next-generation sequencing of brain tissue from the sacrificed animal. **B.** At the molecular level, the Calling Cards transposase, hyperPiggyBac (hyPB) is linked to a tamoxifen-inducible domain, ERT2, which are knocked into a transcription factor (TF) of interest. With transcription and translation of the TF, the TF-hyPB-ERT2 (TF-iCC) fusion protein is produced in the cytoplasm but unable to enter the nucleus due to the unbound conformation of ERT2. Tamoxifen binding allows nuclear translocation. In the nucleus, hyPB can bind to one of the virally delivered transposon sequences. Upon TF-iCC protein binding to a TF gene target, hyPB integrates the transposon into nearby DNA, creating a permanent record of the transient TF-DNA binding event. If SRTs are used, as the transposon has a strong promoter site, it harnesses transcription machinery to “self-report” its location in the genome via mRNA which can then be sequenced with a transposon-selective RNAseq protocol. Created in Biorender.

We have previously adapted the DNA binding domain of SP1 to AAV CC (Cammack et al., 2020). Sp1 is a ubiquitously active TF that binds to unmethylated CpG sites as a proxy for actively transcribed regions (Li et al., 2004). Here, we engineered a knock-in, which enables tagging the full-length protein and adding the ERT2 domain, circumventing existing size limitations of AAV. Sp1-iCC provides a readout of actively transcribed regions during the tamoxifen delivery window, serving as a recorded proxy for gene expression.

We also sought to test whether CC could be adapted to be activity-dependent as well as drug-dependent, inspired by Targeted Recombination in Active Populations (TRAP) (Guenthner et al., 2013) TRAP has allowed such recording for circuit-level analysis: using a tamoxifen-inducible Cre recombinase whose expression is driven by an immediate early gene (IEG) promoter, such as *Fos* or *Arc*, TRAP allows recombination and expression of reporter proteins, only in the presence of both tamoxifen and neural activation (Guenthner et al., 2013). Thus, we targeted Jun, an immediate early gene and binding partner of Fos at Activator Protein 1 (AP-1) sites (Rauscher et al., 1988). Much like Fos-TRAP, the Jun-iCC construct should require both tamoxifen and neural activity for recording to occur. Thus, Jun-iCC, should tag activated neurons with a fluorescent reporter, providing an orthogonal approach to TRAP. Uniquely, Jun-iCC could also provide a post-mortem readout of AP-1 binding during the specified time window, thus serving as a proxy for neural activity-dependent gene expression.

We found that both constructs were tamoxifen-inducible, and we were able to successfully tag activated neurons with Jun-iCC. However, the number of recoverable insertions in Jun-iCC was low compared to AAV Calling Cards. With Sp1-iCC, we were able to recover many more recorded binding events; however, we observed a significant reduction in viability in the Sp1-iCC knock-in animals. Overall, these two mouse lines demonstrate the potential for temporally controlled recording of transcriptional states *in vivo* and encourage further development of iCC in the mouse genome.

## RESULTS

### Jun inducible Calling Cards is inducible in vitro

First, we engineered a Jun-inducible Calling Cards (Jun-iCC) knock-in N2a cell line for *in vitro* validation. We chose Jun as our initial transcription factor target to complement existing FosTRAP technologies that utilize immediate early genes to capture neural activity across development or in response to behavioral stimuli (Cammack et al., 2020). Into the endogenous mJun locus, we engineered the hyperPiggyBac (hyPB) transposase, a myc tag, and an ERT2 domain (Yusa et al., 2011). The ERT2 domain is a humanized estrogen receptor that sequesters any fusion protein to the cytoplasm, allowing nuclear translocation only in the presence of tamoxifen (TAM) or TAM metabolites (Donocoff et al., 2020; Indra et al., 1999). Thus, the Jun-hyPB-myc-ERT2 fusion protein should only be transcribed following neural activity and in the presence of TAM metabolites, conferring temporal control to activity-dependent hyPB transposase insertions. Junction PCR and Sanger sequencing of Jun-hyPB-myc-ERT2 region confirmed that the insertion was successful and there were no mutations in any of the domains.

To assess whether Jun-iCC recording in the N2a cell line is TAM metabolite-dependent, we designed a series of validation experiments. First, we transfected the BrokenHeart (BH) transposon into Jun-iCC cells, which provides a low-background fluorescent readout of transposase activity (**Supplemental Figure S1A**) (Cammack et al., 2020). Our results demonstrated that hyPB insertions occur only in the presence of 4-hydroxytamoxifen (4-OHT), a TAM metabolite (**Supplemental Figure S1B**), confirming the inducibility of our system. We observed that neural activity, induced by propofol, did not visibly affect insertion rates in the Jun-iCC line, likely because immediate early genes like Jun are more ubiquitously expressed in tumor cell lines than at baseline in the mouse brain (Angel et al., 1988).

We next sought to assess the quantity and positions of the insertions, and confirm the system would work with the tools we had already developed for *in* vivo CC. We have previously adapted the Calling Card system to function *in vivo*, by delivering both the transposase and transposons using Adeno Associated Virus (AAV) vectors and have shown its ability to record across mouse brain development (Cammack et al., 2020; Moudgil et al., 2020; Yen et al., 2023). In mouse brain, CC relies on two components: a hyPB transposase, which can be delivered virally or knocked into the genome, and an engineered self-reporting transposon (SRT), which is delivered virally. Therefore, to prepare for *in vivo* testing of these new mouse lines constructs, we also sought to benchmark our cell knock-ins with SRTs. The SRT contains an elongation factor-1 alpha (EF1α) promoter, which allows it to engage transcription machinery to transcribe mRNA, which will include the SRT sequence and portion of the flanking genomics sequence, until a cryptic polyadenylation site is reached (Moudgil et al., 2020). Thus, the SRT “self-reports” its location in RNA: via next-generation sequencing and alignment of the gene target on the mRNA to known gene loci, the location of TF(-hyPB)-DNA binding events can be determined. SRTs vastly increase the sensitivity for insertion sites compared to DNA-based recovery methods. The SRT additionally contains a tdTomato red fluorescent protein, which allows for immunofluorescent visualization of cells where active CC recording events occurred. Therefore, we next transfected the self-reporting transposon (SRT) into the cell lines to be able to read out Jun-iCC insertions via next-generation sequencing (Moudgil et al., 2020). Via bulk next-generation sequencing and Homer motif enrichment analysis, we show an expected enrichment for the Jun motif from *in vitro* Jun-iCC recordings, suggesting that the hyPB was indeed increasingly targeted to AP-1 binding sites (**Supplemental Figure S1C**). Notably, the quantity of insertions, was lower in all Jun-iCC lines compared to the unfused hyPB positive control, with a 2-fold difference in insertion number by sequencing (262,121 in Jun-iCC lines versus 473,304 insertions in the positive controls), suggesting that the endogenous knock-in approach results in reduced Calling Cards recording efficiency, perhaps because of the fusion of the proteins, or because Jun expression may be more difficult to capture due to its transient nature.

### Jun inducible Calling Cards mice are viable and does not disrupt behavior in vivo

Given promising *in vitro* data, we targeted the same Jun-iCC construct into mouse embryos to generate Jun-iCC founder mice: within the endogenous mJun locus, we engineered a hyPB-myc-ERT2 fusion protein, generating mice that are heterozygous for this mutation (**Figure 2A**). We performed junction PCRs to confirm insertion sites, and Sanger sequencing to verify that no unexpected mutations were introduced (**Figure 2B**). We generated two founder lines, F43 and M8. We successfully bred these lines to the fourth generation with almost 200 offspring overall. Offspring showed expected Mendelian ratios (50% for heterozygous breeding), indicating that the Jun-iCC knock-in did not impact viability (**Figure 2C**). Jun-wildtype (WT) indicates two normal copies of the mJun allele, while Jun-iCC indicates a heterozygous knock-in of Jun-hyPB-myc-ERT2 on one WT mJun allele.

**Figure 2.**
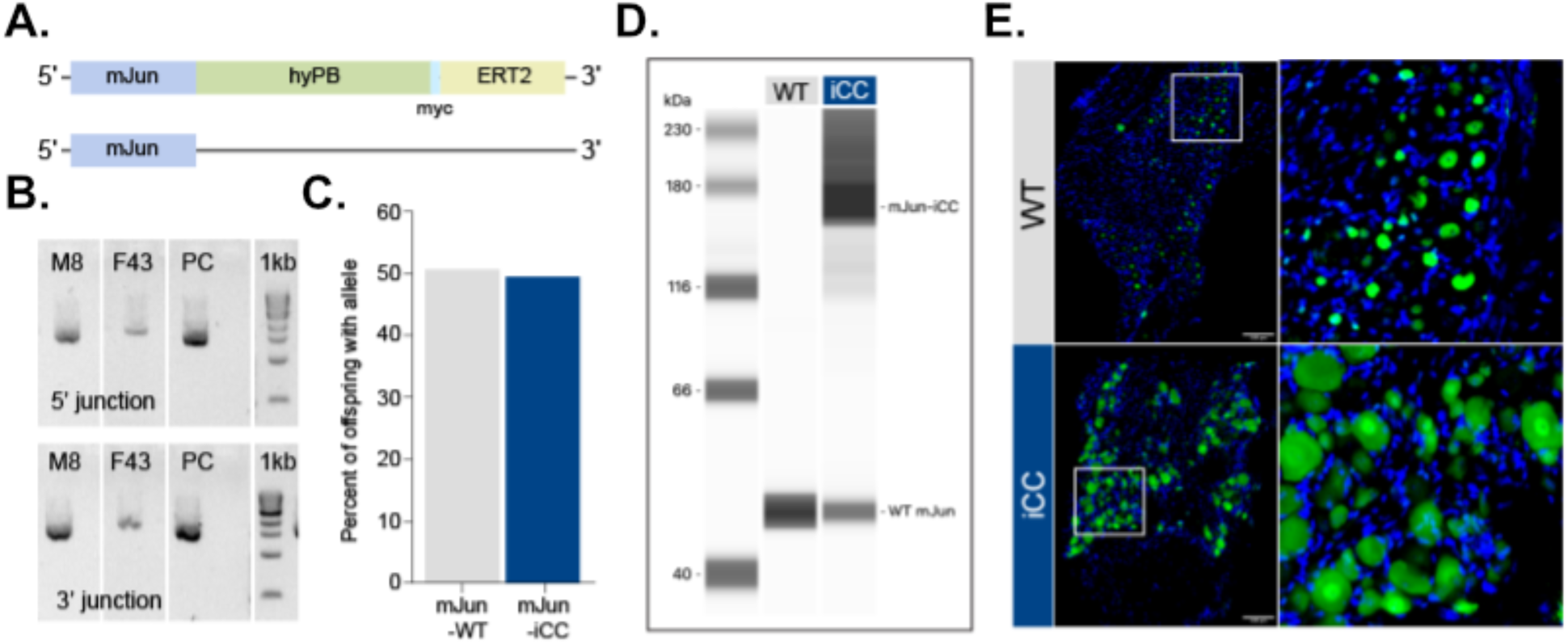
inducible Calling Cards knock-in to mJun is viable. **A.** Schematic of the mJun alleles of the Jun-inducible Calling Cards (Jun-iCC) mouse line. Jun-iCC had one normal copy of mJun, and one copy fused to the Calling Card transposase, hyperPiggyBac (hyPB), a myc tag, and the tamoxifen inducible ERT2 domain. **B.** Junction PCRs showing that both M8 and F43 knock-in lines have expected integration at the mJun locus. **C.** Bar chart showing percent of offspring with either Jun-WT or Jun-iCC allele. Expected inheritance (∼50%) of Jun-iCC allele in N = 179 offspring suggests that Jun-iCC does not impact viability. **D.** Dorsal root ganglia (DRG) samples from Jun-WT mice (with two normal mJun alleles) and Jun-iCC mice (with hyPB-myc-ERT2 knock-in at one mJun allele) were analyzed by Western blot. The wild-type mJun protein (∼40 kDa) is detected in both genotypes, while the larger Jun-iCC fusion protein (∼180 kDa) is observed exclusively in the iCC samples. Molecular weight markers (kDa) are indicated on the left. This confirms expression of the Jun-hyPB-ERT2 fusion protein in the Jun-iCC mouse model. **E**. Immunofluorescence of mJun in DRG sections from Jun-WT (top) and Jun-iCC mice (bottom). Anti-Jun antibody (green) and DAPI nuclear stain (blue). The right panels show enlarged boxed regions. In Jun-WT, Jun localizes primarily to nuclei, while iCC samples show both nuclear and cytoplasmic staining, confirming ERT2-mediated cytoplasmic retention without tamoxifen. Scale: 1000 μm.

Focusing on the M8 transgenic line for the remainder of the work, we confirmed the presence of the full Jun-hyPB-myc-ERT2 fusion protein. Since mJun is only transiently expressed in the brain, we evaluated this in the dorsal root ganglia (DRG), where mJun expression is more durable (Buschmann et al., 1998). We found the wild-type mJun protein (∼40 kDa) in both Jun-iCC and Jun-WT genotypes, while the larger Jun-iCC fusion protein (∼180 kDa) was observed exclusively in the Jun-iCC samples (**Figure 2D**). Immunofluorescence also showed a distinct localization pattern for the Jun-WT versus the Jun-iCC fusion protein, with mJun staining in the Jun-iCC mice displaying both normal nuclear and cytoplasmic localizations (**Figure 2E**). This distinct cytoplasmic localization pattern in Jun-iCC mice confirms that the ERT2 domain is functioning as designed, sequestering the fusion protein in the cytoplasm until tamoxifen administration triggers its translocation to the nucleus, a critical feature for achieving temporal control of the Calling Cards system.

Finally, we tested whether the Jun-iCC mice displayed normal behavior (**Table 1**). In a cohort of forty-four mice, evenly split by genotype, we tested locomotion, sensorimotor skills, anxiety, and olfaction. First, in the open field task, we observed no significant differences in activity levels or exploratory behavior between the two genotypes (**Figure 3A**) (Seibenhener & Wooten, 2015). We found a significant effect of sex, such that males entered the center more frequently and spent longer there than females. Next, in the sensorimotor battery, we found that motor and balance ability were intact in Jun-iCC mice relative to Jun-WT offspring (**Figure 3B**). Of note, Jun-iCC animals exhibited a slight reduction in platform, pole, and 60°-inclined screen tests compared to Jun-WTs. On the platform test, Jun-iCC animals had a lower average time balancing on the platform than Jun-WTs animals, only in batch 1. On the pole test, Jun-iCC animals had a lower average time on the pole than Jun-WTs animals, only in batch 2. On the 60°-inclined screen test, Jun-iCC animals had a higher average time to climb to the top of the screen than Jun-WTs animals, only in batch 2. Notably, Jun-iCC animals performed normally on the 90° inclined screen and inverted screen tests, which are the most arduous tests of balance and strength, suggesting an overall normal phenotype in Jun-iCC mice relative to Jun-WT controls. In terms of sex effects overall, we saw that males had a lower average time balancing on the platform than females, only in batch 1. In the elevated plus maze, we found no differences in time spent in the open arms between genotypes, indicating that Jun-iCC mice do not show increased avoidance (**Figure 3C**) (Walf & Frye, 2007). Jun-iCC animals travelled less distance than Jun-WTs in the EPM, but they displayed normal exploratory behavior and activity in the open field, which is an overall better task to evaluate activity, given its hour-long testing duration and wider, open field. For both open- and closed-arm entries, there was a significant Group × Batch interaction, driven primarily by batch effects, and Jun-WT animals entered the open and closed arms more frequently than Jun-iCC animals only in batch 1. We also found a significant effect of sex, such that males entered the open arms more frequently and spent longer there than females. Finally, in the odorant hole-poking task, we saw that Jun-iCC and Jun-WT mice both were able to distinguish different odors and had comparable numbers of nosepokes for a social odor, suggesting intact olfaction and similar preference for social stimuli (**Figure 3D**)(Wozniak et al., 2013). In summary, we confirmed that a heterozygous iCC construct insertion at the endogenous mJun locus does not robustly impact fundamental behaviors, including locomotion, balance, avoidance, and olfaction.

**Figure 3.**
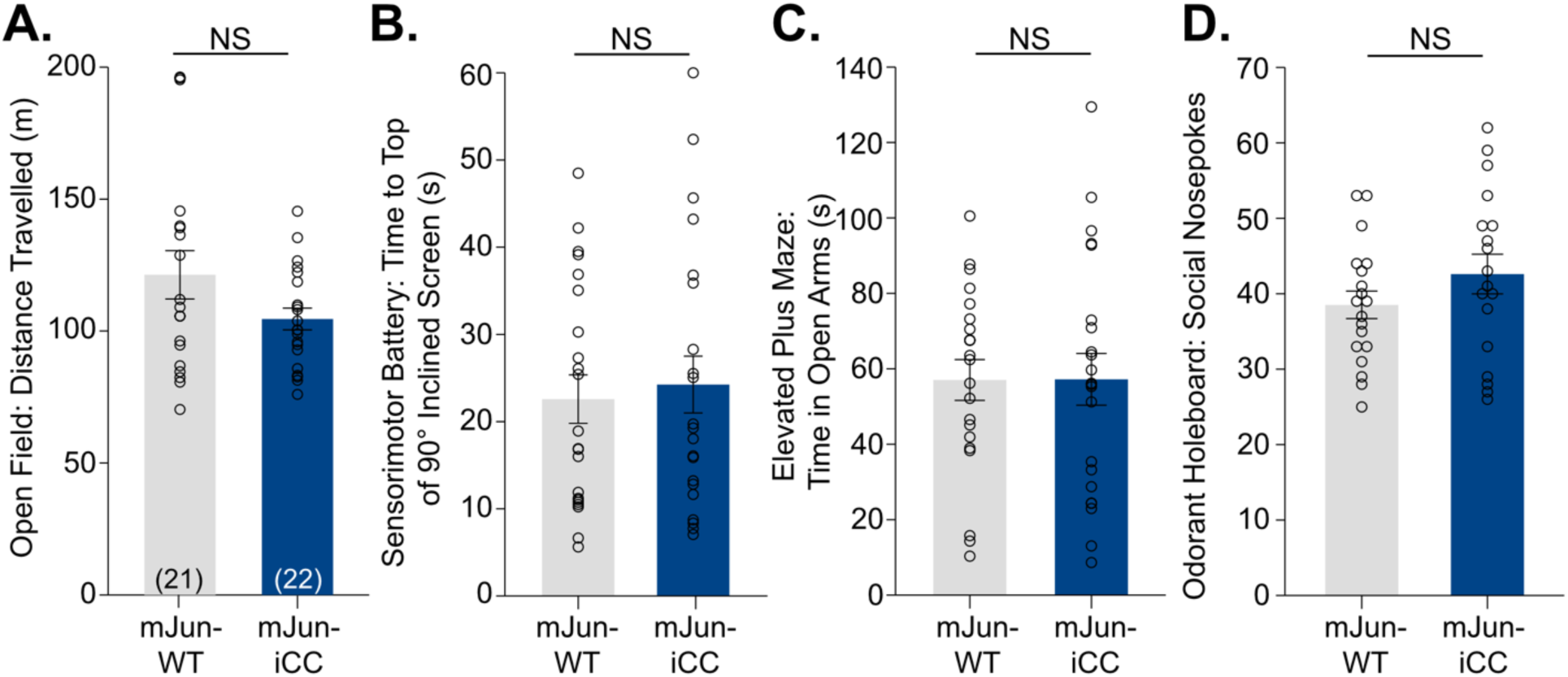
Jun-inducible Calling Card (iCC) mice show comparable behavior to Jun-wildtype (WT) littermates. **A.** No significant differences were observed between Jun-iCC and Jun-WT mice in activity levels in open field. **B.** Motor and balance ability in the sensorimotor battery was also intact in Jun-iCC mice. **C.** Jun-iCC mice did not show increased avoidance in elevated plus maze. **D.** iCC mice showed normal olfactory function and similar preference for social stimuli compared to Jun-WT littermates in the odorant hole-poking task. Bars represent mean ± SEM, individual dots show data from each animal. NS, Not Significant.

**Table 1.** Statistical analysis of outcomes from Jun-iCC behavioral testing. OF, Open Field; SMB, Sensorimotor Battery; HB, Odorant Holeboard; EPM, Elevated Plus Maze.

| Behavioral Assay | Outcome | Model | Predictor | Output | p Value |
| --- | --- | --- | --- | --- | --- |
| OF | Distance | Mann-Whitney | Group | $U(N_{\text{Jun-WT}} = 19, N_{\text{Jun-iCC}} = 21) = 158, z = -1.124$ | $p = 0.261$ |
| | Center Entries | ANOVA | Sex | $F(1,32) = 12.305$ | $p = 0.001$ |
| | Center Time | ANOVA | Sex | $F(1,32) = 8.248$ | $p = 0.007$ |
| SMB | Time on Platform | Mann-Whitney | Batch 1: Group | $U(N_{\text{Jun-WT}} = 10, N_{\text{Jun-iCC}} = 10) = 30, z = -2.163$ | $p = 0.031$ |
| | | | Batch 1: Sex | $U(N_{\text{Jun-WT}} = 13, N_{\text{Jun-iCC}} = 7) = 28, z = -1.984$ | $p = 0.047$ |
| | Time on Pole | Mann-Whitney | Batch 2: Sex | $U(N_{\text{females}} = 11, N_{\text{males}} = 13) = 25, z = -2.694$ | $p = 0.007$ |
| | Time to Top of 60°- screen | Mann-Whitney | Batch 2: Group | $U(N_{\text{Jun-WT}} = 12, N_{\text{Jun-iCC}} = 12) = 36, z = -2.078$ | $p = 0.038$ |
| HB | Social Nosepokes | ANOVA | Group | $F(1, 29) = 0.192$ | $p = 0.665$ |
| HB | Total Ambulation | ANOVA | Group | $F(1, 29) = 2.867$ | $p = 0.101$ |
| EPM | Distance | ANOVA | Group | $F(1,35) = 7.750$ | $p = 0.009$ |
| | | | Group*Sex*Batch | $F(1,35) = 3.370$ | $p = 0.046$ |
| | Open Entries | ANOVA | Sex | $F(1,35) = 6.167$ | $p = .018$ |
| | | | Group*Batch | $F(1,35) = 5.399$ | $p = .009$ |
| | Closed Entries | ANOVA | Group | $F(1,35) = 7.125$ | $p = .011$ |
| | | | Group*Batch | $F(1,35) = 6.122$ | $p = .005$ |
| | Open Time | ANOVA | Sex | $F(1,35) = 7.564$ | $p = .009$ |
| | | | Group*Batch | $F(1,35) = 3.851$ | $p = .031$ |
| | Closed Time | ANOVA | Sex | $F(1,35) = 12.818$ | $p = .001$ |

### Jun-iCC is silent without neural induction in vivo

Next, we benchmarked the Jun-iCC mouse line *in vivo* to validate whether it requires induction by both TAM and neural activity. As a “negative control”, we turned on Jun-iCC recording while animals were in the home cage to test if Jun-iCC is neural activity dependent. At baseline, adult mice have low levels of mJun expression (Ortiz et al., 2020), which should lead to low levels of expression of the Jun-iCC fusion protein, and thus low levels of SRT insertions into the DNA as marked by low RFP fluorescence. If Jun-iCC depended on neural activity for recording, we would expect little detectable levels of RFP following a week of recording in the home cage. We tested four groups: Jun-iCC and WT mice, receiving TAM or vehicle (Veh, cornflower oil and ethanol; *n* = 4). We delivered SRTs via AAV9 intracranially to P1 animals, as before (**Figure 4A**) (Cammack et al., 2020). At P77, we dosed mice with TAM at 100 mg/kg every other day, over the course of five days (Jahn et al., 2018). This regimen leads to TAM presence in the brain for 7 days, meaning Jun-iCC should be “on” for 7 days. A week after the last TAM dose, at P88, we sacrificed animals and harvested their brains for immunofluorescence, focusing on the dentate gyrus, one of the few regions that is active at baseline (Pofahl et al., 2021). We observed no RFP-positive cells in any WT mice or in Jun-iCC mice that received Veh treatment (**Fig 4B**). We saw very low levels of RFP-positive cells in the Jun-iCC mice treated with TAM, only in the dentate gyrus, suggesting that Jun-iCC is silent without neural induction *in vivo*.

**Figure 4.**
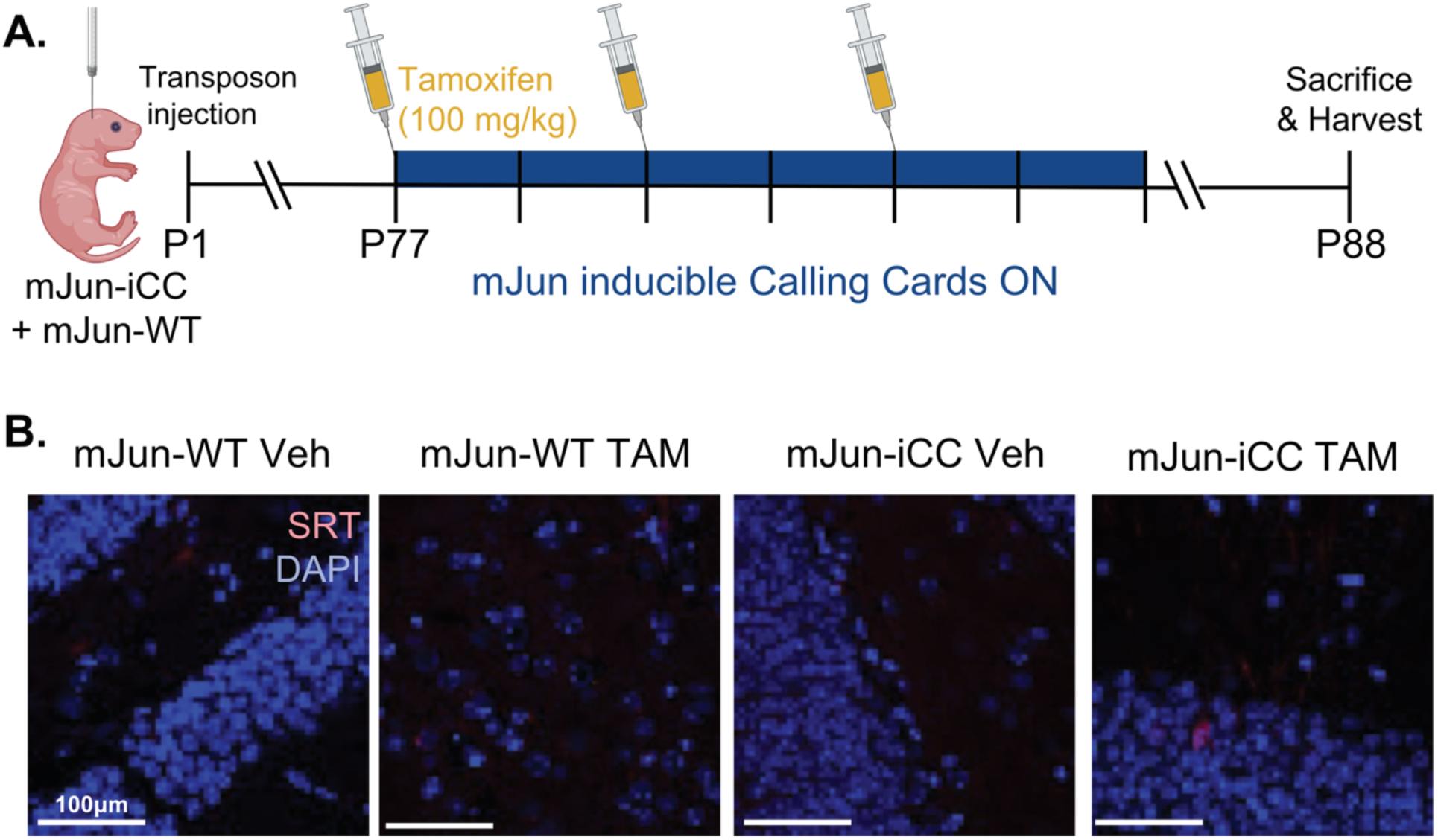
Jun-inducible Calling Cards (Jun-iCC) is silent without neural induction. **A.** Timeline: Transposon was injected into Jun-iCC (iCC) or Jun-WT (WT) pups at postnatal day (P)1. At adulthood (P77), mice were dosed with tamoxifen (TAM) or vehicle (Veh) for 5 days. The dosing scheme should lead to tamoxifen presence for 7 days, thus activating Calling Cards recording for 7 days. Mice were sacrificed and brains harvested for immunofluorescence 7 days after the last TAM dose. **B.** Immunofluorescence shows that transposon levels (as measured by RFP fluorescence) were only present at low levels, in the dentate gyrus of the iCC, TAM-dosed animal, suggesting low levels of baseline Jun-iCC activity in the home cage. Scale bar: 100 uM.

### In vivo Jun-iCC recording depends on TAM and neural activity

Given that mJun levels at baseline in the adult mouse brain are low, we need a “positive control” to validate that Jun-iCC is indeed TAM-*and* activity-dependent. To this end, we induced pharmacological seizures with GABA antagonist pentylenetetrazol (PTZ) while Jun-iCC recording was on (Van Erum et al., 2019). First, we demonstrate that treating with PTZ (30 mg/kg) induced mild seizures in all animals, and latency to seizure behavior was not impacted by sex, genotype, or TAM treatment (**Figure 5A**) (Racine, 1972; Van Erum et al., 2019). We injected both Jun-iCC and Jun-WT mice (*n* = 7) with SRT on P1. In adulthood, mice were administered TAM or Veh, for 5 days, as above. On day 4, we delivered 30 mg/kg PTZ intraperitoneally to all animals (**Figure 5B**) and monitored seizure severity and latency to freezing. All animals exhibiting mild seizures (Racine, 1972; Van Erum et al., 2019), and brains were harvested 8 days after PTZ injection. The left hemisphere of the brain was used for IF for mJun detection, while the right hemisphere was used for RNA sequencing. We found that RFP-positive cells were present only in the dentate gyrus, a region implicated in seizures, in the Jun-iCC TAM animal, with no RFP-positive cells in the other groups (**Figure 5C**) (Erdtmann-Vourliotis et al., 1998; Vasilev et al., 2018). We replicated this experiment in a separate cohort of mice, finding similarly that Jun-iCC, TAM-treated mice had a variable but higher area of RFP+ densities in the dentate gyrus, indicating Jun-iCC recording activity, compared to the Jun-iCC Veh-treated and the Jun-WT TAM-treated controls, which had negligible RFP+ density area (**Supplemental Figure S4**).

**Figure 5.**
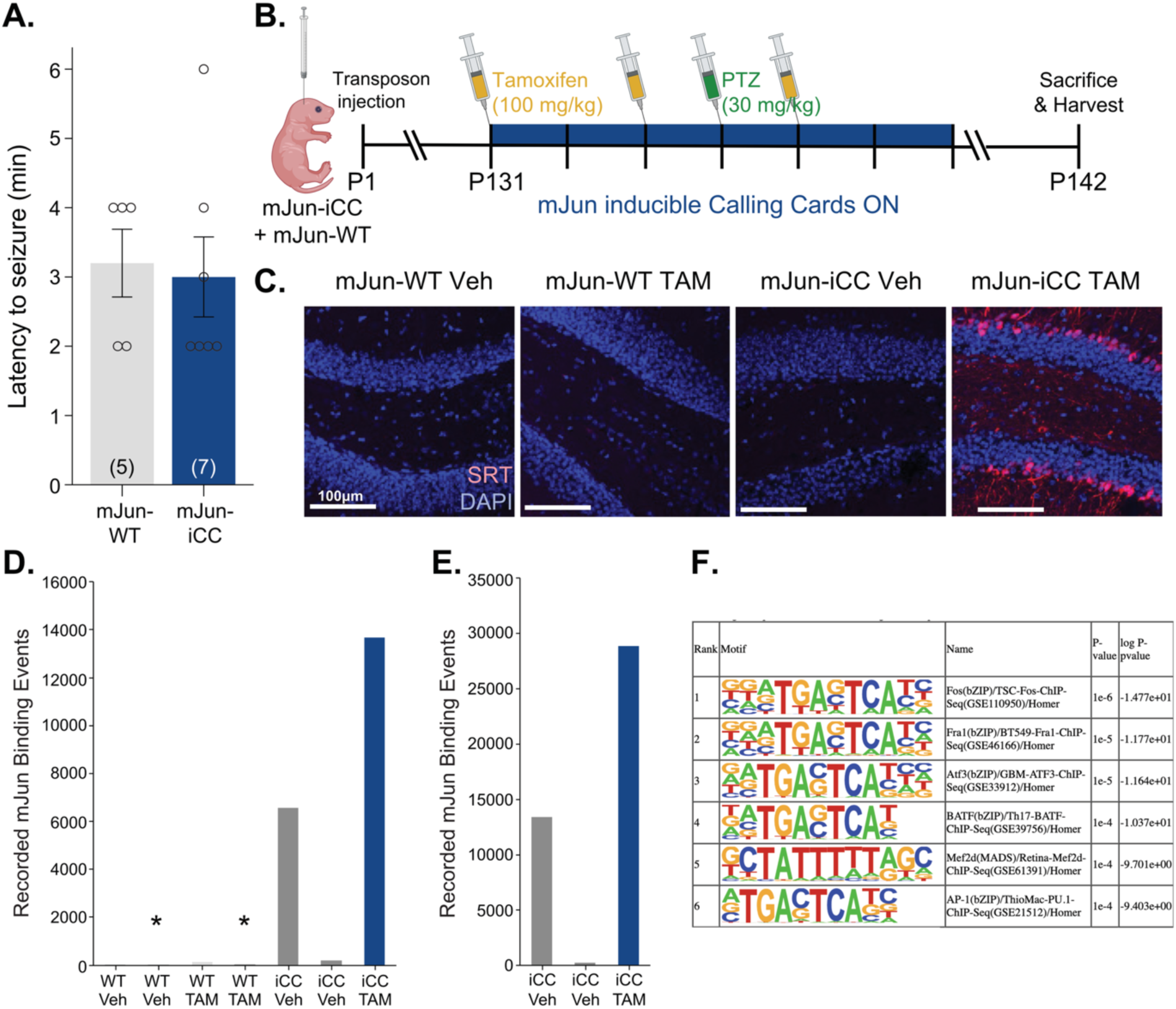
Jun-inducible Calling Cards (Jun-iCC) is induced with tamoxifen and neural activity. **A.** Jun-iCC (iCC) animals do not show increased susceptibility to seizures relative to Jun-wildtype (WT) littermates. **B.** Timeline: Transposon was injected into Jun-iCC (iCC) or Jun-WT (WT) pups at postnatal day (P)1. At adulthood (P131), mice were dosed with tamoxifen (TAM) or vehicle (Veh) for 5 days. The dosing scheme should lead to tamoxifen presence for 7 days, thus activating Calling Cards recording for 7 days. Mice were injected with pentylenetetrazol (PTZ) to induce low-severity seizures on day 4 of TAM-induced Calling Cards recording. Mice were sacrificed and brains harvested for immunofluorescence and next generation sequencing 7 days after the last TAM dose. **C.** Immunofluorescence of dentate gyrus (DG) shows that only the iCC, TAM-dosed animal had RFP-positive neurons, indicating active Calling Cards recording of PTZ-induced seizures. **D.** Next generation sequencing of 1/5^th^ of the brain demonstrated that virtually no mJun binding events were recorded by Calling Cards in the Jun-WT animals regardless of Veh or TAM treatment. Approximately 3x more mJun binding events were recovered from the TAM-dosed animal than the average of the two Veh-dosed animals. **E.** When the entirety of the brain was sequenced for iCC animals only, again approximately 3x more mJun binding events were recovered from TAM-dosed than from Veh-dosed animals. **F.** Homer motif analysis shows significant enrichment for mJun-associated motifs in recovered transposon insertions from sequencing of the entirety of the brain for TAM-dosed iCC animals. Asterisks in bar plots represent samples that did not meet the threshold (>1000 aligned reads).

Next, we performed next-generation CC sequencing to identify the number of recorded mJun binding events. The right hemisphere was divided into 5 samples for technical replicates (Yen et al., 2023). First, we processed only 1/5^th^ of the right hemisphere of all animals. We obtained approximately 3x more SRT insertions from the Jun-iCC TAM-dosed animal compared to the two Jun-iCC Veh-dosed animals (**Figure 5D**, **Table 2**). Then, we more deeply sampled the Jun-iCC Veh and TAM animals and sequenced their entire right hemisphere. Again, we found approximately 3x more recorded mJun binding events in the TAM-dosed relative to the Veh-dosed Jun-iCC mice, with almost 30,000 insertions recorded from the right hemisphere of the Jun-iCC TAM mouse (**Figure 5E**, **Table 3**). Jun-WT mice had virtually no insertions regardless of treatment. Notably, 30,000 insertions per hemisphere is an order of magnitude lower than insertions obtained using viral delivery of non-inducible hyPB (Cammack et al., 2020). This significant reduction in insertions is likely due to the transient nature of mJun expression in the mouse brain, even after a robust stimulus such as an induced seizure. Finally, we performed Homer motif enrichment analysis (Heinz et al., 2010) on the insertions from the Jun-iCC TAM-treated animal, and we found significant enrichment in mJun and related (AP1, Fos) motifs (**Figure 5F**), indicating we were marking promoters likely bound by mJun during the seizure.

**Table 2.**
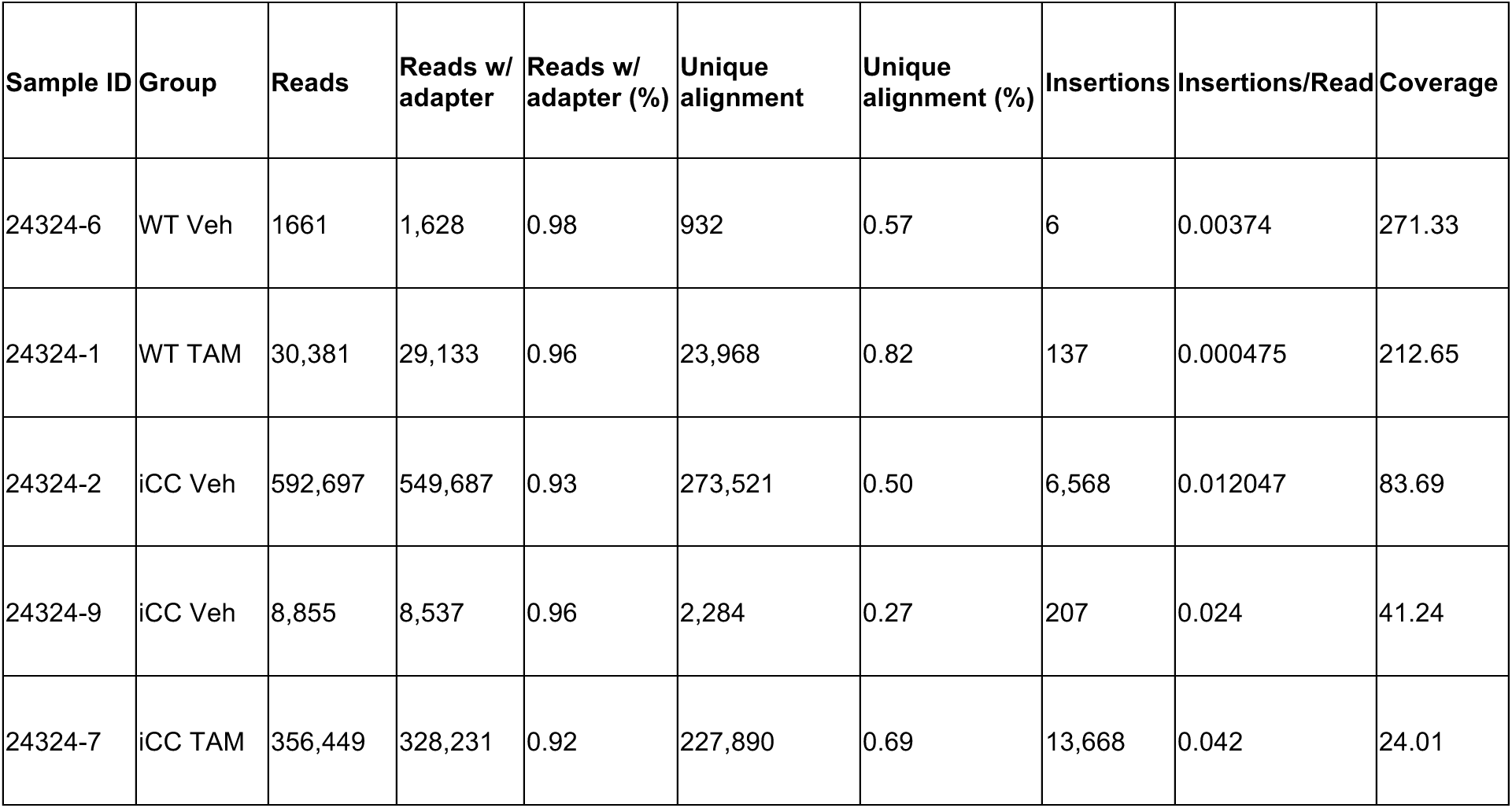
Jun-iCC Sequencing Results table for 1/5^th^ of Brain. Notably, samples with less than 1000 aligned reads were removed.

**Table 3.** Jun-iCC sequencing table for whole brains from positive animals.

| Sample ID | Group | Reads | Reads w/<br>adapter | Reads w/<br>adapter (%) | Unique<br>alignment | Unique<br>alignment (%) | Insertions | Insertions/Read | Coverage |
| --- | --- | --- | --- | --- | --- | --- | --- | --- | --- |
| 24324-2 | iCC Veh | 1,102,982 | 1,038,919 | 0.942 | 633,045 | 0.609 | 13,426 | 0.013 | 77.38 |
| 24324-9 | iCC Veh | 10,794 | 10,430 | 0.966 | 3,497 | 0.335 | 256 | 0.025 | 40.74 |
| 24324-7 | iCC TAM | 819,042 | 773,731 | 0.945 | 529,806 | 0.685 | 28,856 | 0.037 | 26.81 |

### Sp1 inducible Calling Cards is inducible in vitro

Given the relatively low levels of recorded insertions with Jun-iCC, potentially due to its transient expression pattern, we next opted to introduce inducible Calling Cards to a more ubiquitously expressed locus. We chose Sp1 since it is a widely expressed marker of CpG unmethylated regions and thus can serve as a proxy for gene expression (Cammack et al., 2020; Li et al., 2004). Unlike Jun-iCC, which is dependent on both neural activity and tamoxifen for activation, the Sp1-iCC system leverages the constitutive expression of Sp1 across cell types to capture gene expression states more broadly and with higher sensitivity, potentially overcoming the limited recording efficiency we observed with the activity-dependent mJun system.

We engineered an Sp1-inducible Calling Cards (Sp1-iCC) knock-in N2a cell line for *in vitro* validations, with the hyPB-myc-ERT2 knocked into the C-terminus of the endogenous Sp1 locus. Junction PCR and Sanger sequencing of Sp1-hyPB-myc-ERT2 region confirmed that the insertion was successful and there were no mutations in any of the domains. As above, we transfected the BrokenHeart (BH) transposon into Sp1-iCC cells. Using BH, we verified that hyPB insertions happen only in the presence of 4OHT (**Supplemental Figure S3**).

### Sp1 inducible Calling Cards negatively impacts viability

We then knocked in the Sp1-iCC construct into mouse embryos to generate Sp1-iCC founder mice, with hyPB-myc-ERT2 knocked into the endogenous Sp1 locus (**Figure 6A**). We performed junction PCRs to confirm insertion sites and Sanger sequencing to verify no mutations were introduced (**Figure 6B**). We generated two founder lines, M5 and M9. Offspring of both lines did not show expected Mendelian ratios, with only ∼30% of total live offspring being Sp1-iCC heterozygous, compared to the expected 50%, suggesting that the Sp1-iCC knock-in negatively impacts viability (**Figure 6C**). Indeed, when looking at offspring from both founder lines we saw noticeable reductions in litter size from the expected 6-8 pups per litter and increased death of pups immediately after birth or around weaning (**Figure 6D**) (Finlay et al., 2015). We also noticed aberrant phenotypes in the founders and offspring, with anophthalmia being the most prominent. We also saw significant reductions in weight between Sp1-iCC mice and their Sp1-WT littermates (p = 0.005; **Figure 7A**). Overall, this suggests that sequestering one endogenous copy of Sp1 into the cytoplasm negatively impacts viability, may sometimes be incompatible with life, and if offspring survive, negatively impacts their development.

**Figure 6.**
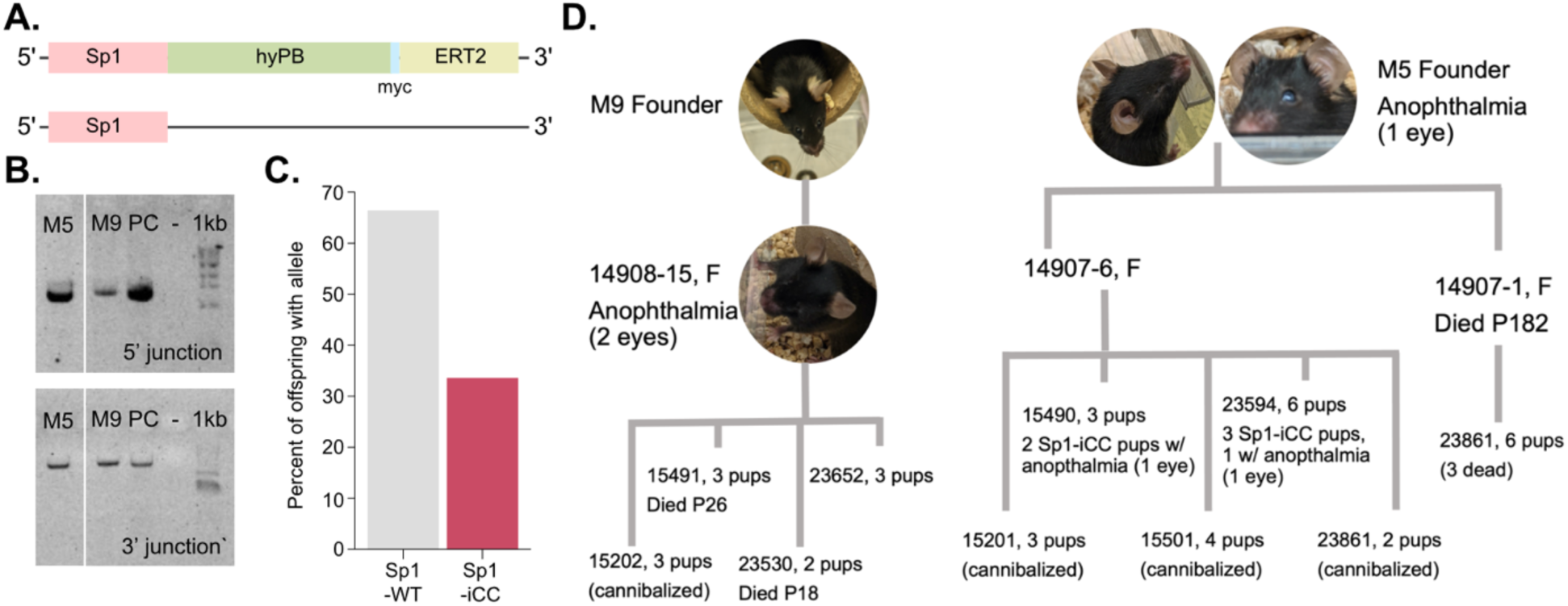
Inducible Calling Cards knock into the Sp1 locus is not a viable strategy. **A**. Schematic of the Sp1 alleles of the Sp1-inducible Calling Cards (Sp1-iCC) mouse line. Sp1-iCC had one normal copy of Sp1, and one copy fused to the Calling Card transposase, hyperPiggyBac (hyPB), a myc tag, and the tamoxifen inducible ERT2 domain. **B.** Junction PCRs showing that both M5 and M9 knock-in lines have expected integration at the Sp1 locus. **C.** Percentage of offspring (N = 131) carrying the Sp1-iCC allele is lower than expected mendelian 50%, potentially indicating reduced viability of Sp1-iCC pups relative to Sp1-wildtype (WT) pups. **D.** Representative family tree of litters from two separate founder lines indicating widespread presence of anophthalmia, small litters, early loss of pups, and mortality around weaning age, overall suggesting limited viability of the Sp1-iCC mouse lines.

**Figure 7.**
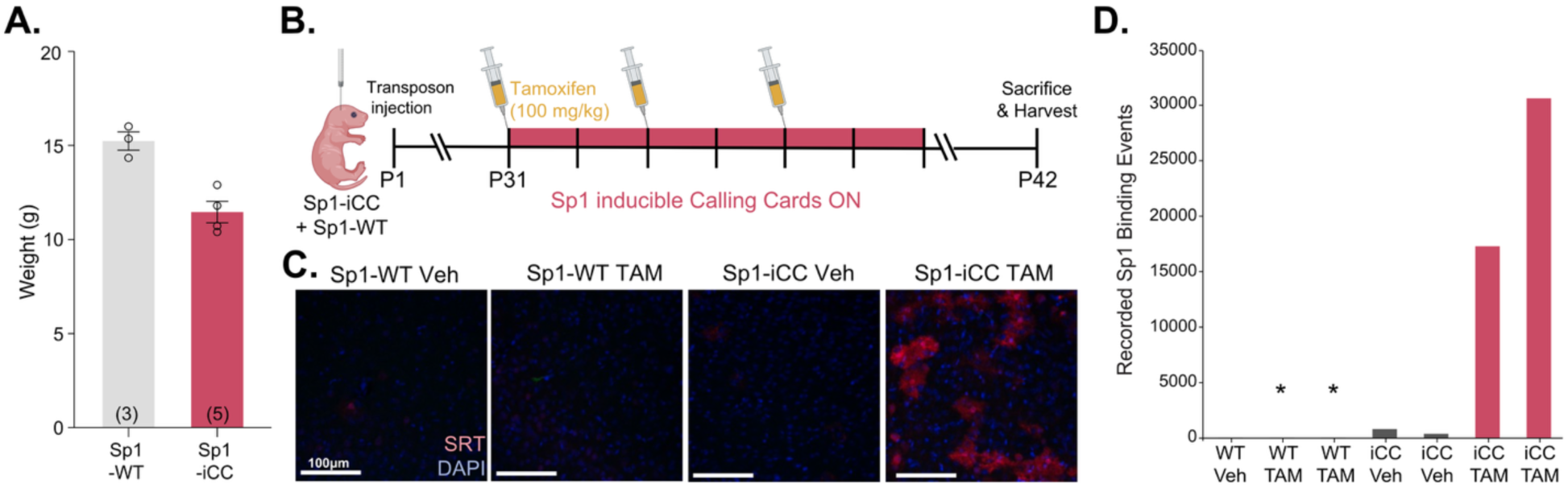
Sp1-inducible Calling Cards (Jun-iCC) is induced with tamoxifen. **A.** Sp1-iCC (iCC) animals weigh less than Sp1-wildtype (WT) littermates (p=0.005). **B.** Timeline: Transposon was injected into Sp1-iCC (iCC) or Sp1-WT (WT) pups at postnatal day (P)1. At juvenile age (P31), mice were dosed with tamoxifen (TAM) or vehicle (Veh) for 5 days. The dosing scheme should lead to tamoxifen presence for 7 days, thus activating Calling Cards recording for 7 days. Mice were sacrificed and brains harvested for immunofluorescence and next-generation sequencing 7 days after the last TAM dose. **C.** Immunofluorescence of cortex shows that only the iCC, TAM-dosed animal had RFP-positive neurons, indicating active Calling Cards recording of Sp1 binding. **D.** Next-generation sequencing of 1/5^th^ of the brain demonstrated that virtually no Sp1 binding events were recorded by Calling Cards in the WT animals, with tens of thousands of insertions recovered from the TAM-dosed animals. Scale bar = 100 uM. Asterisks in bar plots represent samples that did not meet the threshold (>1000 aligned reads).

### In vivo Sp1-iCC recording depends on TAM

To test whether Sp1-iCC recording *in vivo* was TAM-inducible, we tested four groups: Sp1-iCC and WT mice (*n* = 7), receiving TAM or Veh, for 5 days, as with Jun-iCC above (**Figure 7B**). All mice had received SRT injections intracranially on P1. We sacrificed animals a week after their last dose and harvested brains, with the left hemisphere for IF and the right hemisphere for sequencing. We found that RFP-positive cells, including neurons and astrocytes, were only present in the cortex of Sp1-iCC TAM mice, with no RFP-positive cells in the other groups (**Figure 7C**). We replicated this experiment in a separate cohort of mice, finding similarly that only the Sp1-iCC, TAM-treated mice had a noticeably higher number of RFP+ neurons in cortex as well as dentate gyrus, with negligible levels of RFP+ neurons in the brains of control littermates (**Supplemental Figure S4**).

Finally, we performed next-generation sequencing in the first cohort to identify the number of recorded Sp1 binding events in 1/5^th^ of the right hemisphere of all animals. We obtained no SRT insertions in the Sp1-WT animals and negligible amounts of SRT insertions in the Veh-dosed Sp1-iCC animals. The Sp1-iCC TAM-dosed animals had robust recording of Sp1 binding events in these animals (**Figure 7D**, **Table 4**). Notably we obtained over 30,000 insertions from only a fifth of a hemisphere from Sp1-iCC, compared to approximately 30,000 from a full hemisphere with Jun-iCC, suggesting that Sp1-iCC does express more ubiquitously and thus leads to much more recording. Still, while Sp1-iCC provides superior recording efficiency, the developmental impacts in this mouse line will make it challenging to use for many applications.

**Table 4.** Sp1-iCC Sequencing Table for whole brains. Notably, samples with less than 1000 aligned reads were removed.

| Sample ID | Group | Reads | Reads w/ adapter | Reads w/ adapter (%) | Unique alignment | Unique alignment (%) | Insertions | Insertions/Read | Coverage |
| --- | --- | --- | --- | --- | --- | --- | --- | --- | --- |
| 15490-4 | WT Veh | 18,619 | 1,662 | 8.93 | 55 | 3.31 | 24 | 0.014 | 69.25 |
| 15490-7 | iCC Veh | 8,050 | 1,867 | 23.19 | 1,578 | 84.52 | 814 | 0.44 | 2.29 |
| 15490-3 | iCC Veh | 11,505 | 1,043 | 9.066 | 931 | 89.26 | 373 | 0.36 | 2.80 |
| 15490-2 | iCC TAM | 101,731 | 79,111 | 77.76 | 72,289 | 91.38 | 17,320 | 0.22 | 4.57 |
| 15490-6 | iCC TAM | 108,159 | 89,729 | 82.96 | 79,803 | 88.94 | 30,709 | 0.34 | 2.92 |

## DISCUSSION

Our study presents the development and characterization of inducible Calling Cards (iCC) technology, a novel approach for temporally controlled recording of TF-DNA binding events *in vivo*. We engineered two distinct mouse lines targeting different transcription factors: Jun-iCC and Sp1-iCC, each revealing important biological constraints and potential applications for molecular recording in neuroscience.

The Jun-iCC system successfully demonstrated tamoxifen-dependent inducibility both *in vitro* and *in vivo*, with low background recording in the absence of tamoxifen or neural activity. These mice showed normal development and behavior across multiple domains, including locomotion, sensorimotor function, anxiety-related behaviors, and olfaction. When activated by tamoxifen during PTZ-induced seizures, Jun-iCC effectively tagged activated neurons with RFP and recorded approximately 30,000 insertions per hemisphere. The recorded insertions are lower than expected for non-activity-dependent CC tools(Boros et al., 2025; Cammack et al., 2020; Yen et al., 2023), likely due to the transient expression pattern of immediate early genes in the brain. While this recording efficiency may be sufficient for TRAP-like approaches to mapping circuit activation, it appears insufficient for comprehensive epigenetic profiling of Jun binding unless tenfold more mice were used (as hundreds of thousands of insertions would be required to facilitate peak calling).

In contrast, the Sp1-iCC system provided substantially higher recording efficiency, yielding over 30,000 insertions from just one-fifth of a hemisphere. This confirms our hypothesis that targeting a ubiquitously expressed transcription factor would enhance recording capabilities. However, this came at a significant cost to viability. Sp1-iCC mice exhibited reduced Mendelian ratios (∼30% instead of the expected 50%), developmental abnormalities including anophthalmia, reduced body weight, and high mortality rates among offspring.

Interestingly, the severe developmental impact of Sp1-iCC suggests that Sp1 heterozygosity may be functionally lethal or severely detrimental, which aligns with previous literature on Sp1 heterozygosity, where heterozygous mice also display anophthalmia in one or both eyes and growth restriction, as well as decreased erythroid progenitor cells (Krüger et al., 2007; Marin et al., 1997). However, our added sub-viability suggests that Sp1-iCC may even have some dominant negative activity that makes it more severe than heterozygous loss of function, perhaps sequestering key Sp1 binding partners outside the nucleus. Our findings provide further insight into Sp1’s essential role in development. In contrast, Jun-iCC mice exhibited normal development and behavior, consistent with observations from other IEG-inducible models such as Fos-TRAP and Arc-TRAP (Guenthner et al., 2013). This strengthens the suggestion that a single functional copy of immediate early genes is sufficient for normal development, potentially due to redundancy in their gene function.

The Jun-iCC system demonstrates that IEGs derived from transposons can effectively mark activated neurons with fluorescent proteins, akin to Fos-TRAP. Detection of fluorescence requires relatively few copies of tdTomato mRNA to visualize a cell, making it suitable for cellular labeling. However, comprehensive transcriptional recording requires hundreds of thousands of insertions per mouse to be applicable for downstream analysis, which is orders of magnitude more than what is currently achievable with transiently expressed factors like Jun (Yen et al., 2023). Nevertheless, this approach might still prove valuable in tissues where Jun expression is more sustained, such as dorsal root ganglia, where recording efficiency could reach levels suitable for molecular analysis (Buschmann et al., 1998). Likewise, the current iteration may be sufficient to provide an orthogonal or complementary approach to TRAP, but with reporters and other transgenes delivered by transposon rather than LoxP-mediated recombination.

For future iterations of inducible Calling Cards technology, the most promising approach would be to target a non-endogenous locus, such as the Rosa26 locus, with either a ubiquitous transcription factor fused to hyPB or the native hyPB, which has natural affinity for BRD4 at super enhancer sites (Cammack et al., 2020; Chu et al., 2016; Moudgil et al., 2020). Such a construct would avoid the developmental consequences of disrupting essential endogenous genes while maintaining temporal control through the ERT2 domain. In this scenario, using just the Sp1 promoter region or first exon to drive expression and knocking this construct into the Rosa26 locus, rather than modifying the endogenous Sp1 locus itself, could provide the efficiency benefits of Sp1-driven expression without the detrimental impacts on viability.

A currently untestable question in behavioral neuroscience is why genetically identical, similarly reared littermate mice dichotomize into susceptible versus resilient groups in various models, including chronic social defeat stress (CSDS), addiction, and social motivation (Krishnan et al., 2007; Maloney et al., 2023; Slivicki et al., 2023). In the well-studied CSDS model, resilient and susceptible groups demonstrate significantly different transcriptional profiles at sacrifice (Krishnan et al., 2007). Hence, the leading hypothesis is that resilience is predicated by differing molecular landscape from susceptibility mice prior to the stressor (Nestler & Waxman, 2020). To test this hypothesis, a tool to precisely record transcriptional state in a limited window prior to behavioral manipulation and then stop recording during the behavioral manipulation would be ideal. Thus, further updates of the iCC system *in vivo* would be of interest.

Overall, our development of inducible Calling Cards technology successfully demonstrates the potential for temporally controlled recording of transcriptional states *in vivo*, while highlighting important biological constraints when implementing such systems with specific transcription factors, such as Jun and Sp1. These mice are no longer live for further current experimentation, but frozen sperm has been nominated to Jackson labs/MNMRC for distribution to other labs interested in using this current generation of iCC tools. Future iterations should consider utilizing the endogenous hyPB with its natural affinity for BRD4 to capture super enhancer sites, or alternative ubiquitous transcription factors delivered exogenously rather than knocked into endogenous loci, which may avoid the developmental consequences of sequestering essential genes.

## METHODS

### In vitro validation of inducible Calling Cards

To validate tamoxifen-dependent transposase activity, we transfected Jun-hyPB-ERT2 knock-in N2a cell lines with BrokenHeart (BH) reporter plasmid using Neon electroporation (1050 V, 30 ms, 3 pulses). Cells were treated with 1 μM 4-hydroxy tamoxifen (4OHT) or vehicle control, and 16.8 μM propofol was used to induce mJun expression 24 hours post-transfection (Kidambi et al., 2009). tdTomato fluorescence was assessed to confirm transposase activity.

For insertion site analysis, we transfected Jun-iCC cells with self-reporting transposon (SRT-PURO), treated with 4OHT, and selected with puromycin (3 μg/ml). Wild-type N2a cells co-transfected with SRT-PURO and unfused hyPB served as controls. Propofol (16.8 μM) was added before cell harvest to induce mJun expression. Genomic DNA was extracted, and insertion sites were identified by next-generation sequencing, as below. We recovered 262,121 unique insertion events from Jun-iCC cells compared to 473,304 from controls. Insertion site specificity was analyzed using Homer motif enrichment analysis (Heinz et al., 2010).

Similar validation experiments were performed with Sp1-hyPB-ERT2 knock-in N2a lines using the BH reporter system to confirm 4OHT-dependent transposition without propofol stimulation, as Sp1 activation does not require additional stimulus. In brief, two Sp1-iCC cell lines were transfected with BH as above and treated with 1 μM 4OHT or vehicle control. tdTomato fluorescence was again assessed to confirm transposase activity.

### Generation of inducible Calling Cards mouse lines

CRISPR-Cas9 mediated homology-directed repair was used to generate transcription factor (TF)-hyPB-myc-ERT2 knock-in mice. For Jun and Sp1 targeting, guide RNAs were designed and evaluated for off-target effects, with optimal candidates selected based on proximity to target sites (5’ CGTTTTGAGAACAGACTGTCNGG 3’). Donor templates contained left and right homology arms flanking the hyPBase-myc-ERT2 cassette, with Sp1 constructs including an 18aa linker sequence (5’ CAAGTGGGCGGCGGCGCGCCCCGCCTGGGCGGCGGACCCAAGC 3’). Junction PCR primers verified correct integration: for Jun, 5’ junction (F: CTCATACCAGTTCGCACAGGC; R: GGATGTGCTCGTCGTCCAGG) and 3’ junction (F: CTGCGGGCTCTACTTCATCG; R: GGCCCATCTCTCTTGTGACTCTG) yielded 1612 bp and 1675 bp products respectively; for Sp1, 5’ junction (F: CTCATACCAGTTCGCACAGGC; R: GGATGTGCTCGTCGTCCAGG) and 3’ junction (F: CTGCGGGCTCTACTTCATCG; R: GGCCCATCTCTCTTGTGACTCTG) yielded 1360 bp and 1656 bp products respectively. Following validation in N2A cells, fertilized oocytes were electroporated with verified gRNAs, Cas9 protein, and donor templates. Random integration was assessed with TF-specific primers for both 5’ and 3’ integration sites: for Jun, 5’ (F: AACCTTGAAAGCGCAAAACTCCGA; R: GGATGTGCTCGTCGTCCAGG) and 3’ (F: CTGCGGGCTCTACTTCATCG; R: CACTGGGGGCGCCGTCAGGTC); for Sp1, 5’ (F: CAGAACAAGAAGGGAGGCCC; R: GGATGTGCTCGTCGTCCAGG) and 3’ (F: CTGCGGGCTCTACTTCATCG; R: AGTGACATTGGGTGCCACAA). PCR conditions employed SuperFi polymerase with GC enhancer, with specific cycling parameters optimized for each junction. For Jun targets, cycling conditions were: 98°C for 2 min, 37 cycles of 98°C for 20s, 64°C for 30s, 72°C for 2 min, followed by 72°C for 5 min. For Sp1 targets, 5’ junction PCR used SuperFi at 65°C with GC enhancer, while 3’ junction PCR used SuperFi at 64°C with a 69°C/3:00 extension. Successful founders with targeted integration at both junctions were identified by PCR and Sanger sequencing, including Jun-targeted males M2, M8, M34, M45 and females F1, F28, F30, and Sp1-targeted males M5 and M9. Each line was validated for viability and fertility to choose the final candidate line. For both lines, the sperm from two proven male breeders was cryopreserved and stored in two separate locations for security

### Animals

All animal studies were approved by and performed in accordance with the guidelines of the Animal Care and Use Committee of Washington University in Saint Louis, School of Medicine and conform to NIH guidelines of the care and use of laboratory animals. The animals were housed in controlled environments with a 12-hour light-dark cycle, constant temperature and relative humidity, and *ad libitum* access to food and water. Heterozygous males for Sp1-iCC and Jun-iCC were crossed to our in-house C57BL/6J females for breeding. The transgenic lines were refreshed every 8-10 generations by backcrossing to freshly obtained C57BL/6J males and females from Jackson Laboratories (Strain No. 000664). Upon weaning at P21, the animals were group-housed by sex, genotype, and treatment, where applicable. Mouse lines will be available through Jackson Laboratory or the MMRC at stock number (in process).

### Genotyping

Genotyping Tissue (tail biopsy, ear punch, or toe clipping) was obtained from each animal and placed in a PCR tube. 100 µl lysis buffer (25mM NaOH, 0.2mM EDTA, pH 12) was added to each tube and incubated at 99°C for 60 min in a thermocycler. Once the samples cooled to room temperature, 100 µl 40 mM Tris-HCl pH 5 was added to neutralize the alkaline lysis buffer. The crude lysate containing genomic DNA (gDNA) was stored at 4°C. All genotyping PCRs were multiplexed with β-actin_For (AGAGGGAAATCGTGCGTGAC) and β-actin_Rev (CAATAGTGATGACCTGGCCGT) primers, as this not only confirms the presence of gDNA, but also minimizes non-specific amplification. Common hyPB primers were used for both Jun-iCC and Sp1-iCC genotyping (F: TGATGACCTGCAGCAGAAAG; R: GCTGATGTTGTCCCTCAGGT). For each reaction, 1ul crude gDNA was mixed with 5 µl OneTaq Quick-Load 2X Master Mix (New England Biolabs M0271), 1 µl 10µM hyPB For/Rev primer mix, 0.25 µl 10µM β-actin For/Rev primer mix, 2 uL Betaine (ThermoFisher J77507.UCR) and 0.75 µl ddH2O. PCR products were run on a 1% agarose gel and visualized with GelRed (Biotium 41003).

### Protein Expression Analysis of Jun-iCC Fusion Protein

To visualize nuclear translocation of Jun-hyPB-ERT2 (iCC), we used Jun-iCC and Jun-WT littermates. Animals (n=3 per group) received a single dose of tamoxifen (TAM, 100 mg/kg) or vehicle control via intraperitoneal injection. Dorsal root ganglia (DRGs) were harvested 12 hours post-injection. Proteins were extracted using RIPA buffer supplemented with protease inhibitors, quantified by BCA assay, and resolved on 4-15% gradient SDS-PAGE gels. Western blotting was performed using antibodies against c-Jun (rabbit anti-c-Jun 60A8, Cell Signaling Technology 9165, at 1:50) to detect both wild-type mJun (∼40 kDa) and the Jun-iCC fusion protein (∼180 kDa, calculated molecular weight 144 kDa). The presence of bands at expected molecular weights confirmed expression of both proteins in the appropriate genotypes, with the Jun-iCC fusion protein detected exclusively in the iCC samples.

### Open Field

Behavioral assays to evaluate effects of Jun-iCC knock-in on animal behavior were conducted in N = 44 animals, split into two batches (Batch 1 *n* = 20, Batch 2 *n* = 24). Both batches were split by genotype (*N_Jun-iCC_ = 22, N_Jun-WT_ = 22)* and sex (*N_female_ = 24, N_male_ = 20*). All tasks were run during the light phase, by a female experimenter blinded to the groups tested.

The 1-hour Open Field (OF) test was used to assess the general activity, exploratory behavior, and emotionality of the mice (*N_total_ = 44, N_batch 1_ = 20, N_batch 2_ = 24, N_female_ = 24, N_male_ = 20, N_Jun-iCC_ = 22, N_Jun-WT_ = 22).* The task was administered at P47 for Batch 1 (range P44-51) and P62 for Batch 2 (range P60-64). We performed the protocol similarly to our published work (Chaturvedi et al., 2024) The apparatus consisted of matte white acrylic enclosures (40 x 40 x 30cm high) enclosed within a white sound-attenuating chamber (70.5 × 50.5 × 60 cm), with red 9 lux illumination (LED Color-Changing Flex Ribbon Lights, Commercial Electric). Animals were habituated to the testing room for 30-60 minutes before testing. Each animal was placed in the center of the maze and allowed to explore freely for 1 hour. Behavior was recorded using ANY-maze (Stoelting Co.), which established a 28.3 x 28.3 cm central zone (50% of total area) and a bordering 5.9 cm peripheral zone, which are delineated on the computer program only. Outcomes included number of entries into and time spent in center as well as the distance traveled in each zone. The apparatus was cleaned with 0.02% chlorhexidine solution (Nolvasan, Zoetis) between animals. 1 animal from Batch 1 and 3 animals from Batch 2 were excluded from open field analyses due to jumping.

### Sensorimotor Battery

Balance, strength, coordination, and motoric initiation were assessed by a battery of sensorimotor measures (SMB) using the protocols described in Maloney et al. (2018) and Maloney et al. (2019). The series of tasks was administered at P49 for Batch 1 (range P46-53) and P69 for Batch 2 (range P67-71). The battery included walking initiation, ledge, platform, pole, and inclined, and inverted screen tests. For each test, the experimenter manually recorded with a stopwatch. Two trials were conducted for each test, back-to-back, and the average was used for the analyses. Tests are divided into two days and are not counterbalanced to provide each animal with the same testing conditions. The first day of testing consisted of the walking initiation, ledge, platform, and pole tests. The second day of testing consisted of the inclined and inverted screen tests. Each test was conducted for a maximum of 60 seconds, except for the pole test, which lasted for a maximum of 120 seconds. Animals were habituated to the testing room for 30-60 minutes before testing.

Walking initiation was evaluated by placing the mouse on a flat surface inside a square measuring 24 x 24 cm, and recording the time it takes the mouse to leave the square (i.e., all four limbs have crossed the square at the same time). The ledge and platform tests were used to assess basic balance ability. The ledge test requires the mouse to be placed on a Plexiglas ledge measuring 0.5 cm deep and standing 38 cm high. The time the mouse is able to balance on the ledge is recorded. During the platform test, the mouse is placed on a wooden platform measuring 3.5cm thick and 3.0 cm in diameter and elevated 25.5 cm above the base. The time the mouse was able to balance on the platform was recorded. Fine motor coordination was elevated by the pole test, where the mouse is placed head upward on a vertical pole with a finely textured surface. The time the mouse takes to turn downward 180° and climb to the bottom of the pole was recorded. The 60°, 90°, and inverted screen tests are used as measures of strength and coordination. During the inclined screen tests, the mouse is placed head oriented downward in the middle of a mesh wire grid measuring 16 squares per 10 cm, elevated 47 cm, and inclined to 60° or 90°. The time the mouse spends turning upward 180° and climbing to the top of the screen is recorded. For the inverted screen test, the mouse is placed head oriented downward in the middle of a mesh wire grid measuring 16 squares per 10 cm, elevated 47 cm, and when it is determined that the mouse has a proper grip on the screen, it is inverted to 180°. The time the mouse is able to hold on to the screen without falling off is recorded. The apparatuses were cleaned with a 70% ethanol spray between animals, and between trials if excrement was in the way.

### Elevated Plus Maze

The elevated plus maze (EPM) was used to assess anxiety-like behavior by measuring avoidance of open spaces. The task was administered at P83 for Batch 1 (range P80-87), and P94 for Batch 2 (range P92-96). We performed the protocol as described previously (Nygaard et al., 2023). The apparatus (Kinder Scientific) consisted of two open arms and two closed arms arranged in a plus configuration and elevated above the floor. Animals were habituated to an adjacent room for 30-60 minutes prior to testing. Testing was conducted in darkness with each animal placed in the center of the maze and allowed to explore freely for 5 minutes. Behavior was recorded using Ethovision software (Noldus). Measures included number of entries into and time spent in open arms, closed arms, and center zone, as well as distance traveled in each zone. The apparatus was cleaned with 0.02% chlorhexidine (Nolvasan, Zoetis) solution between animals.

### Odorant Holeboard

The holeboard (HB) exploration/olfactory preference task was used to evaluate olfaction, social preferences, and exploratory behavior. The task was administered at P51 for Batch 1 (range P48-55) and P73 for Batch 2 (range P71-75). We used the protocol adapted from Maloney et al. (2019). The apparatus is a computerized holeboard (41 x 41 x 38.5 cm) with eight equidistant holes in the floor (Learning Holeboard; MotorMonitor, Kinder Scientific, LLC, Poway, California). Beam breaks quantified the frequency and duration of holepokes (2 cm deep) and dips (1 cm deep). Animals were habituated to the testing room for 30-60 minutes before testing. Each animal completes the test over two days. On the first day, each mouse was given a 30-minute habituation session during which the holes contained no odorants. The following day, a 20-minute test session was conducted with three corner holes baited, as shown in Figure X, with (1) a familiar odorant (corncob bedding soiled with urine and scent markings from same-sex mice), (2) a novel odorant (corncob bedding soiled with urine and scent markings from opposite-sex mice), and (3) a novel, putatively rewarding odorant (vanilla extract diluted onto a filter). The odorants were contained in a cup at the bottom of the hole (7 cm deep) and access was blocked by a plastic mesh cap. The configuration of the odorant-containing and empty corner holes was counterbalanced within and across groups. Exploration was quantified by measuring total ambulation. Olfactory preference was assessed by comparing holepoke frequencies between odorant-containing and empty corner holes. Specifically, we focused on comparing social holepoke frequencies between the same-sex bedding odorant and the opposite-sex bedding odorant. All odorant-containing cups were cleaned with mild soap and water, and the entire apparatus was cleaned with a 0.02% chlorhexidine (Nolvasan, Zoetis) solution. During batch 1 testing, a malfunction in one of the three apparatuses prevented the recording of habituation and test session data for seven mice.

### Statistical Analysis

Behavioral statistical analyses were performed using the IBM SPSS Statistics software (v.28). Prior to analysis, data were screened for missing values and checked for ANOVA assumptions within levels of Genotype, Sex, Batch, and their interactions. Normality was assessed using Shapiro–Wilk tests on z-score–transformed data, along with histograms and Q– Q plots. Levene’s test evaluated the homogeneity of variances. Outliers were identified via boxplots and z-scores exceeding ±3.00. When assumptions were violated, nonparametric Mann– Whitney U tests replaced ANOVAs. Probability value for all analyses was *p* < .05.

### Intracerebroventricular injections

Injections were performed as described in the Intracerebroventricular Injection section within Basic Protocol 1 found in (Yen et al., 2023). Briefly, the pups were anesthetized on ice and a total of 6 µl (3 µl per hemisphere, 1 µl per site) was injected into the ventricles of P0-1 pups using a 50 µl Hamilton syringe. After the injections, the pups were kept warm on a heating pad until they were returned to their home cage.

### Tamoxifen administration

For temporal control of the inducible Calling Cards system, tamoxifen (TAM, Sigma-Aldrich T5648) was administered via intraperitoneal injection. The tamoxifen working solution (10 mg/ml) was prepared by dissolving 50 mg TAM in 5 ml of vehicle solution consisting of sunflower oil (Sigma-Aldrich) and 100% ethanol at a 9:1 ratio. The mixture was vortexed for 30-60 minutes with parafilm-sealed tubes until completely dissolved. Vehicle control solution (Veh) consisted of the sunflower oil and ethanol mixture without tamoxifen. Fresh working solutions were prepared weekly and stored at 4°C.

For induction experiments, mice were weighed prior to the first dose and administered 100 mg/kg tamoxifen (or equivalent volume of vehicle) daily for 5 consecutive days, alternating sides of the intraperitoneal cavity to minimize irritation. All injections were performed using 23-gauge needles to accommodate the viscous solution. TAM-treated and Veh-treated animals were housed separately to prevent cross-contamination through coprophagia. For all experiments, animals were sacrificed at least 7 days after the final tamoxifen dose to allow for clearance of the drug and completion of the recording window.

### Immunofluorescence and imaging

Animals were deeply anesthetized with Isoflurane in an induction chamber until unresponsive to toe pinch. The right hemisphere was processed as per *Bulk Calling Cards library* preparations (Yen et al., 2023). The left hemisphere was harvested and drop-fixed in a tube containing 4% (w/v) PFA overnight at 4°C. Then the tissue was cryoprotected in 15% (w/v) sucrose, then 30% (w/v) sucrose at 4°C, then frozen in plastic molds (Polysciences 18646A-1) containing OCT compound (Fisher Scientific 23-730-571). The tissue blocks were kept at -80°C until further processing. Tissue was cut into 35 µm-thick sagittal or coronal free-floating sections for immunostaining. The sections were permeabilized with 0.1% (v/v) Triton X-100 for 15 mins and blocked with 5% (v/v) normal donkey serum (Jackson ImmunoResearch 014-000-121) for 60 mins. The primary antibodies used were rabbit anti-RFP (Rockland 600-401-379) at 1:500 dilution and rabbit anti-c-Jun (60A8; Cell Signaling Technology 9165). Secondary antibodies used was donkey anti-rabbit Alexa Fluor568 (ThermoFisher A10042) and donkey anti-rabbit Alexa Fluor488 (ThermoFisher R37118). 1 µg/ml DAPI (ThermoFisher D1306) was used to stain the nuclei blue. Sections were mounted onto slides with Prolong Gold anti-fade mounting medium (ThermoFisher P36934) and sealed with nail polish. High magnification confocal images were captured using 20x or 40x objectives on the LSM700 AxioImager Z2 (Zeiss). For quantifying RFP+ area in pixels (Figures S2 and S4), at least two mice per condition and one section per mouse were analyzed.

### Bulk Calling Cards library preparations

Tissue homogenization, RNA isolation, and library preparation steps are described in Basic Protocol 2 and 3 found in (Yen et al., 2023). Briefly, the dissected brain tissue was cut into 5 chunks to identify up to 5 independent insertion events at any given insertion locus, snap-frozen in the vapor phase of liquid nitrogen, then stored at -80°C until further processing. For homogenization, the tissue chunk homogenized in Trizol Reagent (ThermoFisher 15596018) and total RNA was harvested using the RNA Clean & Concentrator Kit-25 (Zymo Research R1018) with slight modifications as described in (Yen et al., 2023). Bulk sequencing libraries were generated and sequenced on the Illumina platform. Calling Cards found at the same insertion site were considered distinct if they had distinct barcodes (i.e. came from different tissue chunks). Insertions that pass filtering were treated equally during analysis, regardless of read depth.

### Sequencing

Pooled dual indexed libraries were submitted to the Genome Technology Access Center at the McDonnell Genome Institute (GTAC@MGI) for sequencing. For their workflow, the concentration of each library was determined using the KAPA Library Quantification Kit according to the manufacturer’s protocol. Target sequencing depth was determined prior to pooling and samples were pooled in ratios based on the targeted depth and concentrations to produce cluster counts appropriate for the Illumina NovaSeq 6000 instrument. Normalized libraries were sequenced on a NovaSeq 6000 S4 Flow Cell using the XP workflow and a 151x10x10x151 sequencing recipe according to the manufacturer’s protocol. Base calls were converted to fastq format and demultiplexed using the onboard DRAGEN software to run BCL Convert.

### Bulk Calling Cards analysis

The raw FASTQ files were processed using the nf-core/callingcards pipeline (Guo et al., 2024; Yen et al., 2023).The resulting qbed files were filtered to keep only insertions with more than 2 reads. Motif enrichment analysis was used from the HOMER suite of tools (Heinz et al., 2010). Common genome arithmetic operations such as merging, intersecting, and counting genome regions were performed using the Bedtools utilities (Quinlan & Hall, 2010).

### Pentylenetetrazol administration and seizure monitoring

To induce neural activity for in vivo testing of the inducible Calling Cards system, we administered pentylenetetrazol (PTZ, a GABA antagonist) to mice (Van Erum et al., 2019). PTZ solution (5 mg/ml) was prepared fresh on the day of use by dissolving 50 mg PTZ in 10 ml of sterile 1X PBS. The mixture was vortexed and incubated in a 37°C water bath for approximately 10 minutes to ensure complete dissolution.

Mice were weighed immediately before injection to calculate the appropriate dose volume. For all experiments, we used a 30 mg/kg dose, calculated as: (0.030 × body weight in grams × 1000)/5 = injection volume in μl. PTZ was administered via intraperitoneal injection, and animals were immediately placed in a clean observation cage with fresh bedding for seizure monitoring.

Seizure activity was scored according to the modified Racine scale (Racine, 1972; Van Erum et al., 2019). For our experiments, we recorded the latency to freezing and monitored animals for a minimum of 15 minutes. All animals displayed mild seizures that did not progress to more severe stages. Following seizure induction, animals were returned to their home cages after regaining normal locomotion and monitored daily for any adverse effects.

## Data and code availability

Raw data, processed data, and code used to analyze the data are available upon request.

## Acknowledgements

We thank members of the Dougherty and Mitra laboratories for helpful discussions and feedback; M. Wallace and the Mouse Genetics Core for their services and expertise in generating the transgenic mice; M. Li and the Hope Center Viral Vectors Core at Washington University School of Medicine for generating the AAV vectors; the DNA Sequencing and Innovation Lab (DSIL) and the Genome Technology Access Center at the McDonnell Genome Institute (GTAC@MGI) for their sequencing expertise and services.

This work was funded by grants from the National Institute of Mental Health (RF1MH117070, RF1MH126723, and R01MH124808 to J.D.D. and R.D.M.). A.Y. was supported in part by T32HG000045 from the National Human Genome Research Institute. S.S. was supported in part by the Autism Science Foundation (22-007).

## Author contributions

S.S., R.D.M., and J.D.D designed research. S.S. A.V., T.A., A.C., X.C., M.P performed research. X.C. and R.D.M contributed unpublished reagents/analytic tools. S.S, A.V, T.A., and A.C. analyzed data. S.S., A.V., and T.A. prepared the figures and data visualization. S.S. A.V., T.A., M.C.C., R.D.M., and J.D.D. wrote the paper. S.S., R.D.M., and J.D.D coordinated the project. S.S., M.C.C., R.D.M., and J.D.D. acquired financial support for the project.

## Conflict of interest statements

R.D.M. has submitted patents related to calling card technology.

## SUPPLEMENTAL FIGURES

**Supplemental Figure S1.**
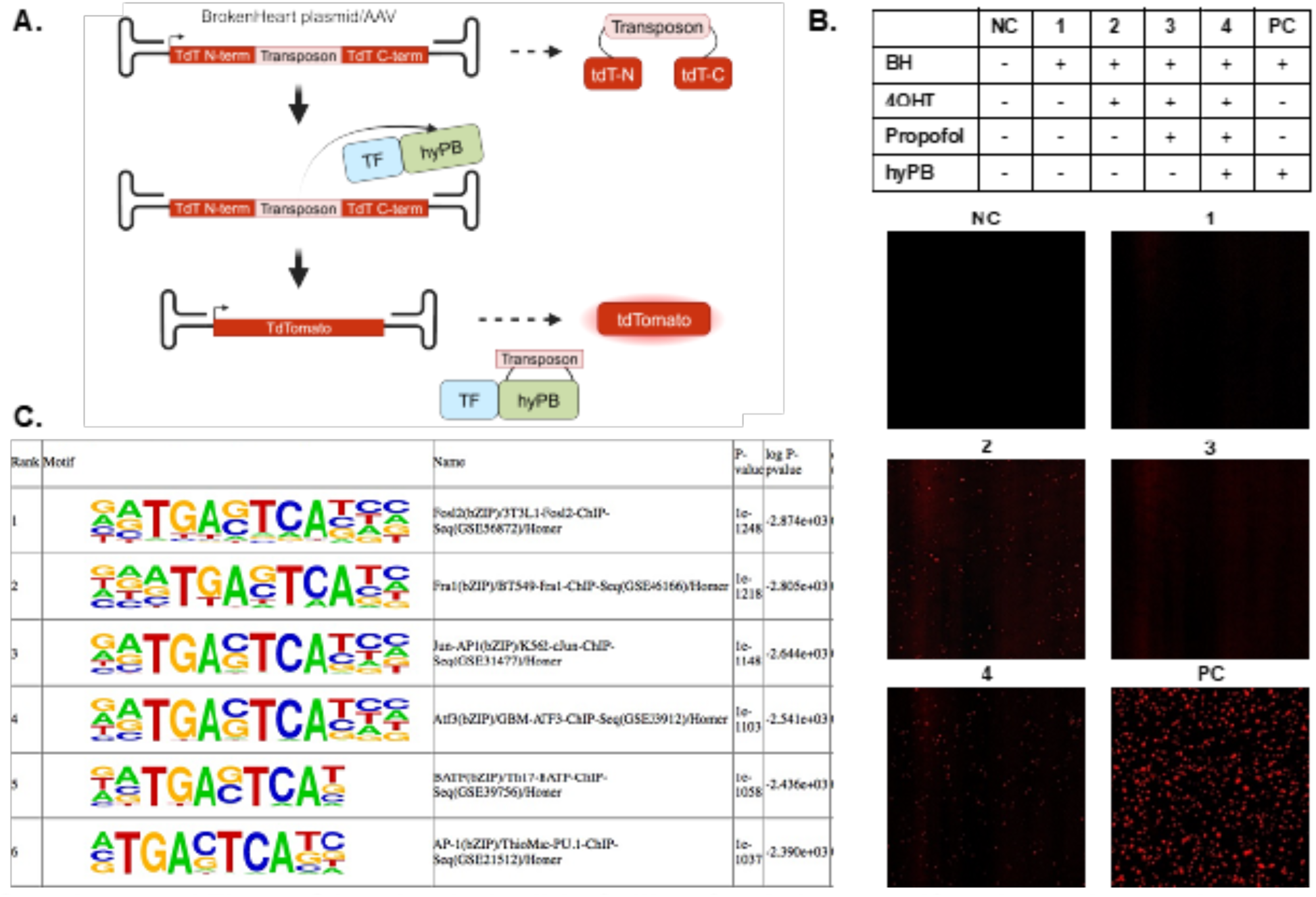
mJun inducible Calling Cards is tamoxifen-dependent and targets expected loci *in vitro*. **A.** Schematic of how BrokenHeart plasmid/AAV acts as a measure of Calling Card transposase, hyperPiggyBac (hyPB) activity. In brief, BrokenHeart sequence contains a transposon in between the N- and C-terminals of tdTomato fluorescent reporter. Transcription of native BrokenHeart leads to non-functional protein, and no red fluorescence. Only in the presence of hyPB activity, with excision of the transposon sequence, can the full tdTomato be transcribed and translated, leading to red fluorescence. **B.** Knock-in lines of Jun-inducible Calling Cards show transposon activity, as measured by tdTomato fluorescence from the BrokenHeart (BH) transposon, only in the presence of tamoxifen metabolite (4OHT), however at a much lower level than the unfused hyperPiggyBac (hyPB) transposase (positive control). **C.** Homer motif analysis shows significant enrichment for mJun-associated motifs in transposon insertions from Jun-iCC knock-in lines.

**Supplemental Figure S2.**
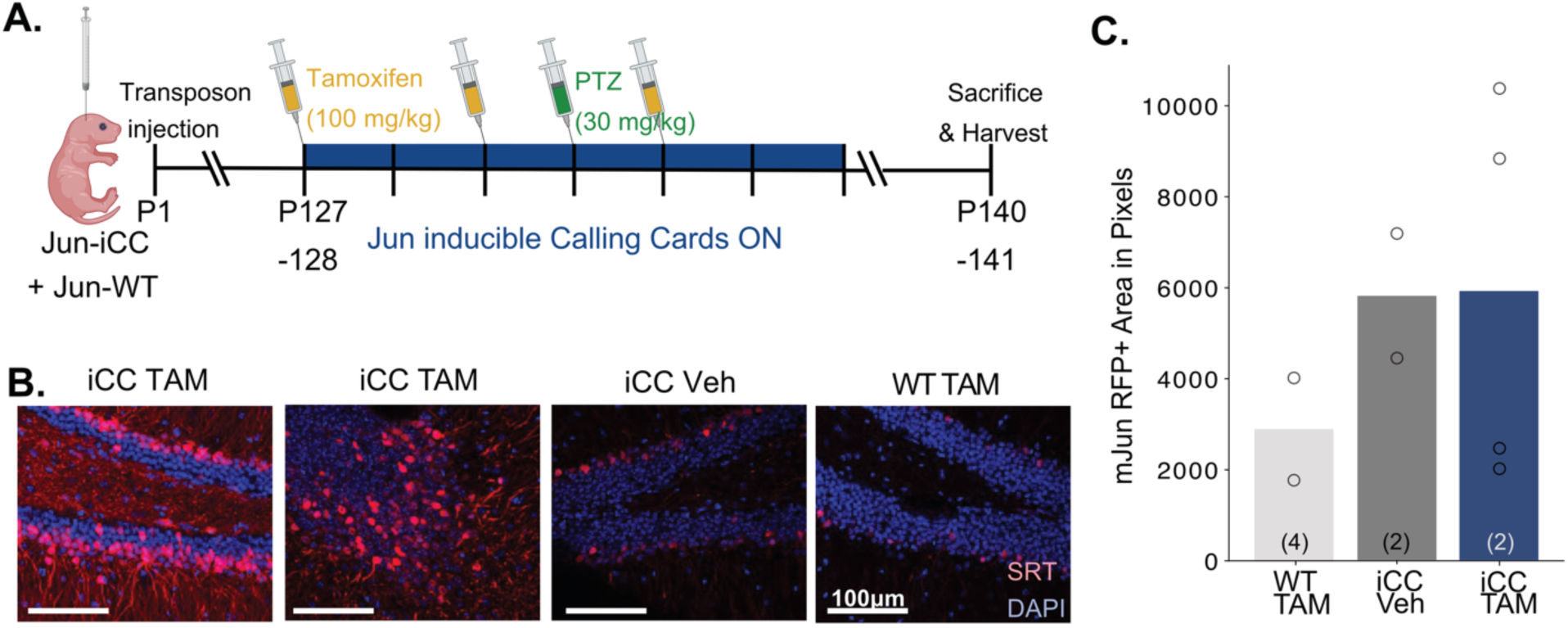
Replication cohort demonstrates Jun-inducible Calling Cards (Jun-iCC) is induced with tamoxifen and neural activity. **A.** Timeline: Transposon was injected into Jun-iCC (iCC) or Jun-WT (WT) pups at postnatal day (P)1. At adulthood (P127/8), mice were dosed with tamoxifen (TAM) or vehicle (Veh) for 5 days. The dosing scheme should lead to tamoxifen presence for 7 days, thus activating Calling Cards recording for 7 days. Mice were injected with pentylenetetrazol (PTZ) to induce low-severity seizures on day 4 of TAM-induced Calling Cards recording. Mice were sacrificed and brains harvested for immunofluorescence 7 days after the last TAM dose, at P140-1. **B.** Immunofluorescence of dentate gyrus (DG) shows that only the two iCC, TAM-dosed animals had RFP-positive neurons, indicating active Calling Cards recording of PTZ-induced seizures. Scale bar = 100 uM. These results from an independent cohort confirm our findings in Figure 5, demonstrating the reproducibility of tamoxifen and activity-dependent recording with the Jun-iCC system across multiple experiments. **C.** RFP+ area within the whole image **(B)** measured in pixels shows that some Jun-iCC animals dosed with TAM had larger RFP+ areas.

**Supplemental Figure S3.**
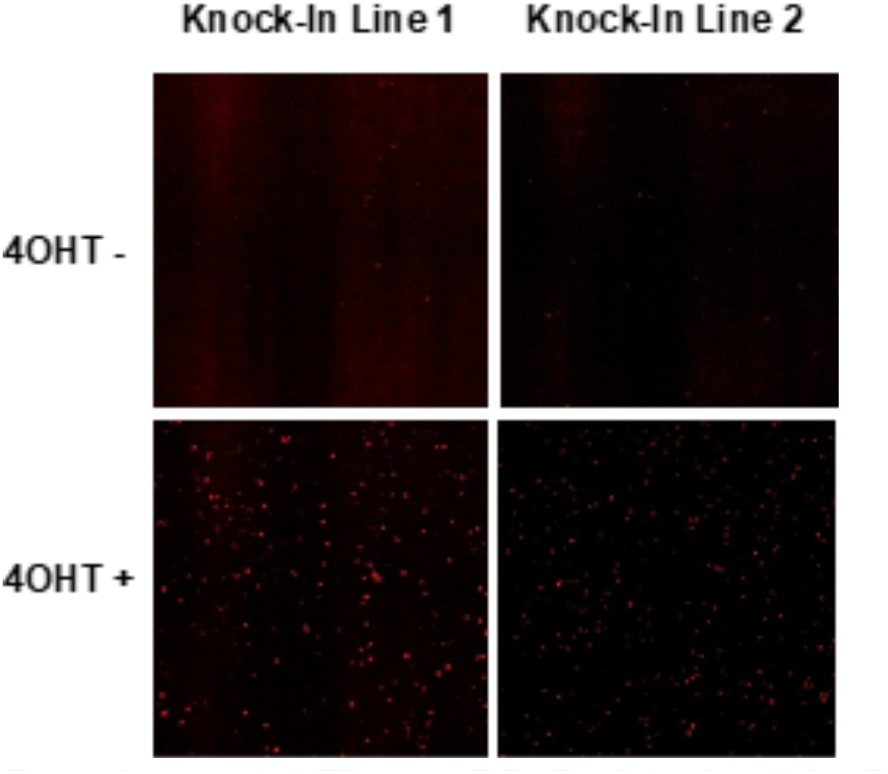
Sp1 inducible Calling Cards is tamoxifen-dependent *in vitro*. Two separate knock-in lines of Sp1 inducible Calling Cards show transposon activity, as measured by tdTomato fluorescence from the BrokenHeart (BH) transposon, only in the presence of tamoxifen metabolite (4OHT).

**Supplemental Figure S4.**
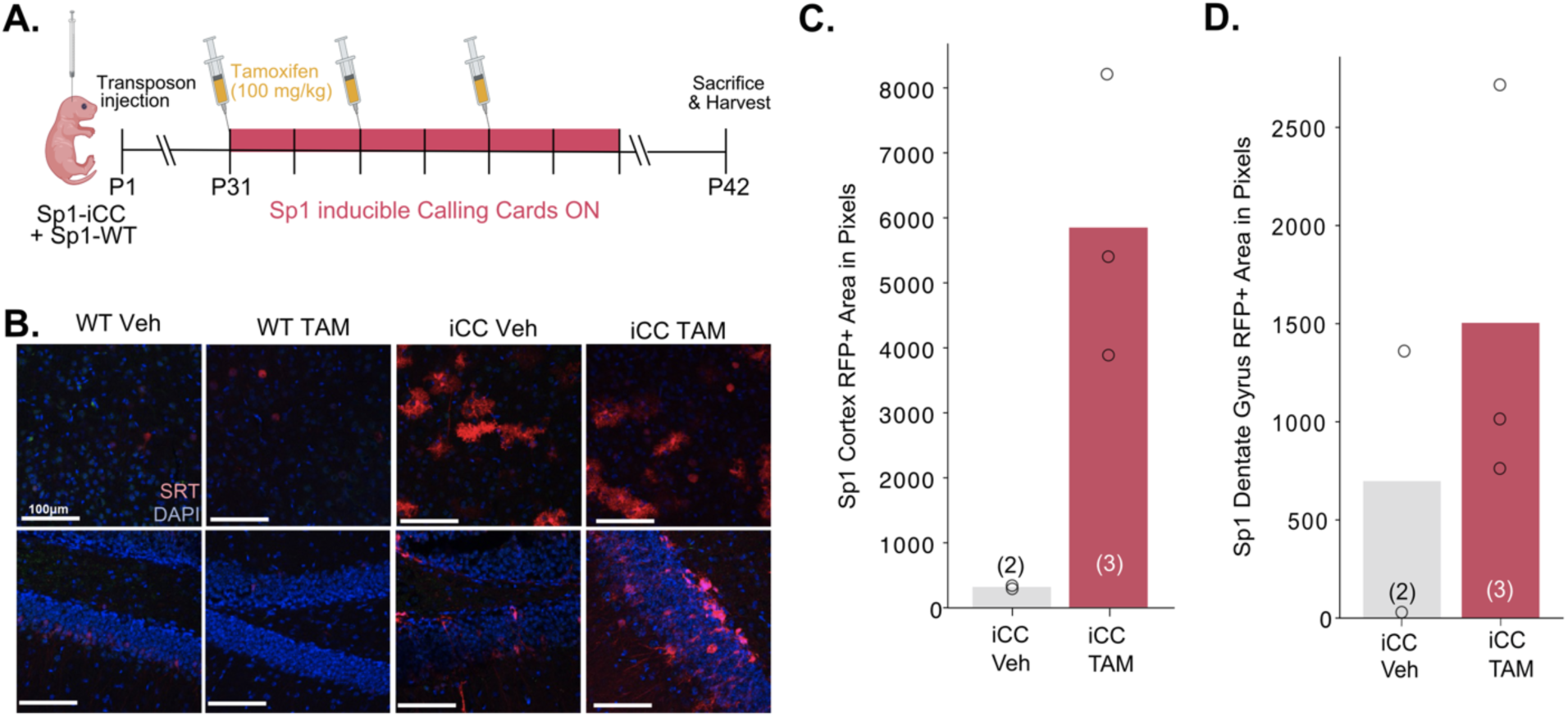
Replication cohort demonstrates Sp1-inducible Calling Cards (Sp1-iCC) is induced with tamoxifen. **A.** Timeline: Transposon was injected into Sp1-iCC (iCC) or Sp1-WT (WT) pups at postnatal day (P)1. At juvenile age (P31), mice were dosed with tamoxifen (TAM) or vehicle (Veh) for 5 days. The dosing scheme should lead to tamoxifen presence for 7 days, thus activating Calling Cards recording for 7 days. Mice were sacrificed and brains harvested for immunofluorescence 7 days after the last TAM dose. **B.** Immunofluorescence of cortex (top panels) and dentate gyrus (bottom panels) shows that only the iCC, TAM-dosed animals had RFP-positive neurons, indicating active Calling Cards recording of PTZ-induced seizures. **(C & D)** RFP+ area within the whole image **(B)** measured in pixels shows that some Jun-iCC animals dosed with TAM had larger RFP+ areas within the cortex and dentate gyrus, respectively.

